# IFI207, a young and fast-evolving antiviral factor, stabilizes STING

**DOI:** 10.1101/2023.01.19.524411

**Authors:** Eileen A. Moran, Karen Salas-Briceno, Alexya N. Aguilera, Thomas M. Keane, David J. Adams, Jingtao Lilue, Susan R. Ross

## Abstract

Mammalian ALR proteins bind nucleic acids and initiate production of type I interferons or inflammasome assembly, thereby contributing to host innate immunity. *ALR*s are encoded at a single genetic locus. In mice, the *Alr* locus is highly polymorphic at the sequence and copy number level. We suggest that one rapidly evolving member of the *Alr* family, *Ifi207*, was introduced to the *Mus* genome by a recent recombination event. *Ifi207* has a large, distinctive repeat region that differs in sequence and length in different *Mus* strains. We show that IFI207 plays a key role in the STING-mediated response to cGAMP, DNA, and MLV, and that IFI207 controls MLV infection in vivo. Uniquely, IFI207 acts by stabilizing STING protein via its repeat region. Our studies suggest that under the pressure of host-pathogen coevolution, in a dynamic locus such as the *Alr*, recombination between gene family members creates new genes with novel and essential functions that play diverse roles in biological processes.

## INTRODUCTION

Innate immune responses are critical to control viral, bacterial, and fungal infections. The mammalian innate immune system consists of numerous pattern recognition receptors (PRRs) that recognize intracellular nucleic acids and other components of pathogens, termed pathogen-associated molecular patterns (PAMPs) (1). PRR activation initiates intricate signalling cascades which culminate in the production of type I interferons (IFNs), proinflammatory cytokines, and inflammatory cell death. These responses also ensure that the adaptive immune system is activated and subsequently generates a specific response to eliminate pathogens. Multiple PRRs likely evolved so that the innate immune system can discriminate between the many different pathogens and PAMPs it encounters (2-5).

One family of PRRs with evidence of positive selection is the Aim2-like receptor (ALR) family (6, 7). Previous studies suggested that there are 4 human *ALR*s and 13–16 mouse *Alr*s all encoded at a single locus (6-9). Although it is not known what selective pressures caused the diversification of this locus, several ALRs have been studied for their role in innate immunity. For example, human interferon-gamma inducible (IFI) 16 and mouse IFI204 are required for type I IFN and cytokine responses to herpes simplex virus-1 (HSV-1) and vaccinia virus (VACV) DNA (10). Several ALRs also affect transcription; IFI16 and IFI204 function in the nucleus to regulate HIV and HSV-1 transcription, while IFI205 regulates the inflammatory response by controlling apoptosis-associated speck-like molecule containing CARD domain (ASC) transcription (11-13). As cytoplasmic sensors, IFI204 is required for IFN production during *M. bovis* and *F. novicida* infection, and IFI203 and IFI205 play a role in the IFN response to murine leukemia virus (MLV) infection and endogenous retroelement DNA detection, respectively (8, 13-15). These ALRs, as well as the innate immune sensors cyclic GMP-AMP (cGAMP) synthase (cGAS) and dead box helicase 41 (DDX41), mediate production of IFNs and cytokines in a stimulator of interferon genes (STING)-dependent manner (3, 8, 13-19). Activated STING dimerizes and interacts with TANK binding kinase 1 (TBK1), thereby stimulating the dimerization and nuclear translocation of the transcription factors IFN regulatory factor 3 (IRF3) and NF-κB, leading to the production of type I IFNs and cytokines (20-23).

Here we show that the Alr locus in *Mus* species has undergone rapid evolution for at least the past few million years, and that one gene in the locus, *Ifi207* (*PyhinA*), is highly divergent even among closely related inbred mouse strains. We found that IFI207 is unique among the ALRs due to a large repeat region separating the two domains common to all ALRs, the N-terminal pyrin domain (PYD) that facilitates homotypic and heterotypic interactions with PYD-containing and other proteins, and the C-terminal hematopoietic expression, interferon-inducible nature, and nuclear localization (HIN) 200 domain that binds DNA (4, 7, 24). We found that the number of repeats is variable among inbred wild and laboratory strains of mice, ranging in number from 11 to 25. Due to the evidence of pathogen-driven positive selection on the *Alr* locus and the evolutionary diversity of Ifi207, we examined the role of IFI207 in innate immune responses. We demonstrate that IFI207 stabilizes STING protein via its repeat region and that allelic variants with longer repeats stabilize STING more effectively. Furthermore, we show that this stabilization enhances the STING response to nucleic acid ligands and cGAMP and that IFI207 plays a role in controlling MLV infection in vivo.

## RESULTS

### *Ifi207* is a young gene exclusively found in the *Mus* genus

We previously reported that the *Alr* locus is highly variable, with different numbers and complements of *Alr* genes mapping to the locus in three inbred mouse strains, C57BL/6, DBA/2J and 129P2/OlaHsd (8). The locus in the GRCm39 reference genome (strain C57BL/6J) encodes 14 genes (Fig. 1A), whilst strain 129P2/OlaHsd encodes 16, and DBA/2J, 17 genes (8). Many members of the *Alr* family are similar to each other, both in the coding and non-coding regions, and are surrounded by transposon elements. To further study the genome structure of *Alr* locus in house mice, we accessed the long read (whole genome PacBio Sequel / Hifi + Chicago HiC scaffolding) de novo assemblies of 13 inbred house mouse strains and *Mus Spretus* and *Mus Caroli* from the Mouse Genomes Project. As shown in Fig. 1A and 1B, based on gene structure and phylogeny, the *Alr* gene family consists of 9 variant members in house mice and other *Mus* species, and the number of genes in the locus ranges from 10 (PWK/PhJ) to 17 (DBA/2J, AKR/J, CBA/J) (Fig. 1A). In these inbred mouse strains, there is one copy of *Aim2* at the 5’ end of the locus followed by 2-3 copies of *Ifi206* homologues (*Ifi206*, *Ifi213* and *Ifi208* in the GRCm39 reference), 1-2 copies of *Ifi214*-like genes, one copy of *Ifi207*, 1-3 copies of *Ifi202*, 1-4 *Ifi203* homologues, 1-2 *Mnda* members, one copy of *Ifi204* and possibly one copy of *MndaL*. *Ifi208*, *Ifi207* and *Ifi204* are encoded in a stable island in the middle of *Alr* locus, whilst other regions are highly dynamic in the evolutionary history of the house mouse.

**Fig. 1.**
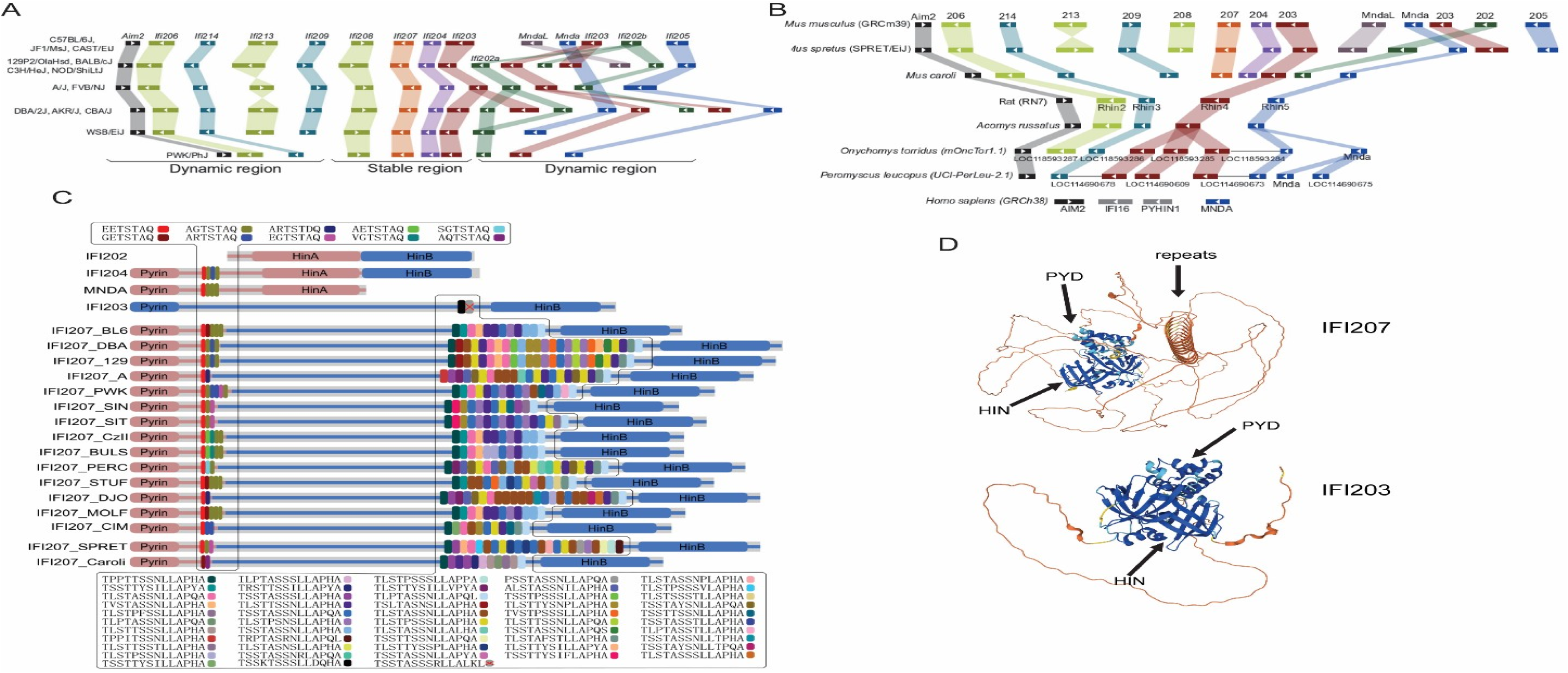
*Ifi207* structure. A) Genome structure of *Alr* locus in inbred mouse strains. Based on gene structure and phylogeny, the 9 family members with variate copy numbers are indicated by the color blocks. *Ifi208*, *Ifi207*, *Ifi204* and one copy of *Ifi203* are encoded in a stable contig, and other regions in the *Alr* locus showed significant numbers of inversions, recombinations and duplications. B) Genome structure of *Alr* locus in other rodent species and humans. *Ifi207*, *MndaL* and *Ifi202* are only found in the *Mus* genus. Some of the genes in Onychomys and Peromyscus were annotated tandemly, probably caused by low annotation quality. C) IFI207 protein structure in different *Mus* genus members. D) Comparison of IFI207 and IFI203 predicted protein structure. Structures were downloaded from AlphaFold (https://alphafold.ebi.ac.uk/) (*52, 53*).

In addition, we downloaded the whole genome assembly from similar long read techniques carried out with other rodent species. When the *Mus* locus is compared with other rodent species, only orthologues of *Aim2*, *Ifi206*, *Ifi214*, *Mnda* and *Ifi203* can clearly be found in the latter (Fig. 1B). Rats have exactly 5 *Alr* members. The Golden spiny mouse (genus *Acomys*), deer mice (genus *Peromyscus*) and grasshopper mice (*Onychomys*; both from family Cricetidae) have the exact same genome structure as rats, except with variable copy numbers of Ifi203 paralogues. No *Ifi207*, *MndaL*, *Ifi204* and *Ifi202* orthologues can be found in these species (Fig. 1B). Furthermore, we compared the de novo assembly of 23 *Euarchontoglires* species and investigated the whole genome Illumina reads of key species in the *Murinae* subfamily. Homologues of *Ifi207* can only be found in the *Mus* genus (Table S1). Indeed, the genome structure around *Ifi207* (including introns) is probably derived from the 3’ end of *Ifi203* and the 5′ end of *Mnda*, from a recent recombination event in the ancestor of the *Mus* genus. Additionally, the 5’ end of *Ifi203* and the 3’ end from *Mnda* possibly fused into *MndaL* in the same recombination event (Fig. S1). In comparison to rodent species, the human genome encodes *AIM2* and *MNDA*, and *IFI16* and *PYHIN1*, the latter with ambiguous relationship to rodent *Alrs* (Fig. 1B) (6).

### *Ifi207* is characterized by two highly polymorphic repeat regions

Most ALRs contain an N-terminal PYD domain and one or two HIN domains, termed HIN-A and HIN-B, depending on their sequence (6, 7). From the de novo assembly of the *Alr* locus in mice, we found two repeat regions in several genes which are highly polymorphic between mouse strains. At around residue 120 (PYD-side repeat) of IFI204, IFI205, MNDA and IFI207 there are 2-6 units of the sequence XXTSTA/DQ. All the repeats are encoded in the same exon, precisely 7 residues as a unit, and the first two residues are highly divergent (A, E, G, R, V, S or Q; Fig. 1C). This repeat was also found in the 2nd exon of *Mnda* in rat and grasshopper and deer mice but no other species, including humans (Fig. S2, Table S2).

The other repeat unit is unique to IFI207 and is located close to the HIN-B domain, with 11 - 25 units of the sequence TXSTXSSXLLAPXA (S/T-rich repeat). In 49 unique repeat sequences extracted from 13 mouse strains, there was up to 36% diversity (average value 15%) between their nucleotide sequence, and up to 65% difference at the amino acid residue level (29% average); this was confirmed with multiple sequencing methods (Fig. 1C and Table S2; see Materials and Methods). Like the PYD-side repeat, all the S/T-rich HIN-side repeats are encoded in a single large exon, with precisely 14 residues (42 base pairs) in a unit. Interestingly, in IFI203, the possible origin and closest paralogue of IFI207, the sequence surrounding the repeat region is conserved between IFI203 and IFI207; however, only one or two copies of the S/T-rich HIN-side repeats are found, and they are highly conserved between mouse strains (Fig. S3). Moreover, a single copy of the same S/T-rich sequence can also be found close to the HIN-B domain of human IFI16 (Fig. S3, Table S2). These results indicate the long S/T-rich repeats in IFI207 were recently introduced and are evolving in a very short time in mouse evolution (see Table S2 for the alignment of Ifi203 and Ifi207). AlphaFold analysis predicts that the ST-rich repeat region forms a coiled structure not found in the other proteins encoded in the *ALR* locus (Fig. 1D). The extremely fast evolution of S/T-rich repeats make *Ifi207* a unique gene in the mouse genome, and indeed, in mammalian species in general.

### Knockout of *Ifi202b*, *Ifi204* or *Ifi207* in mice has a minimal effect on expression of other genes in the locus

To test the function of IFI207, we generated an *Ifi207* knockout (KO) mouse. Mouse *Alr* genes, including both coding and intervening sequences, are highly homologous, making it difficult to precisely delete single genes in the locus8. However, we were able to use CRISPR/Cas9 to delete exon 3 of the *Ifi207* gene in C57BL/6N mice (Fig. S4A). Additionally, mice with floxed alleles of *Ifi202b* and *Ifi204* on a C57BL/6N background were available through the KOMP repository. *Ifi202b* encodes a protein that lacks a PYD and has two HIN domains, A and B, while *Ifi204* encodes a protein with PYD, HIN-A and HIN-B domains (Fig. 1C). The replacement allele in IFI202 KO mice deleted exon 3 and Cre-mediated deletion resulted in IFI204 KO mice lacking exons 3 and 4 (Fig. S4A; see Methods). The knockout alleles for both *Ifi204* and *Ifi207* generate truncated proteins predicted to contain only the PYD, while the *Ifi202* knockout allele generates a protein predicted to contain a portion of the HIN-A domain and a full HIN-B domain (Fig. S4B). All three KO strains had normal viability and produced offspring.

We examined expression of the KO alleles in bone marrow-derived macrophages (BMDMs) and dendritic cells (BMDCs) from wild type and knockout mice using RT-qPCR. *Ifi202* RNA was barely detectable in both cell types from C57BL/6 (BL/6) mice, as we previously reported (8); this was also the case in the KO mice (Fig. 2A). *Ifi204* and *Ifi207* RNAs were not detected in the respective KO strains (Fig. 2A). To determine if *Ifi202*, *Ifi204* or *Ifi207* knockout affected expression of the other genes in the locus, we examined RNA levels in BMDMs and BMDCs from BL/6, IFI202, IFI204 and IFI207 KO mice. BMDMs from STING ^mut^ mice were also included in these analyses. These mice are resistant to stimulation with DNA and other PAMPs as they encode a mutant STING that is unable to signal (25). Moreover, because it has previously been shown that the genes in the Alr locus are type 1 IFN-inducible, we tested whether deletion of *Ifi202b*, *Ifi204* or *Ifi207* affected induction by IFN-β (6, 26, 27). Neither basal nor IFN-β-induced levels of expression of the different genes were altered in either BMDMs or BMDCs by knockout of any of the ALRs or the STING mutation, apart from *Mnda*; *Mnda* RNA levels were consistently reduced in BMDMs and BMDCs in IFI204 KO mice in the absence and presence of IFN-β (Fig. 2A). The significance of this observation is unknown. Nonetheless, since loss of *Ifi207* did not affect the expression of the other *Alrs*, phenotypes observed in the KO mouse are likely specific to this gene.

**Fig. 2.**
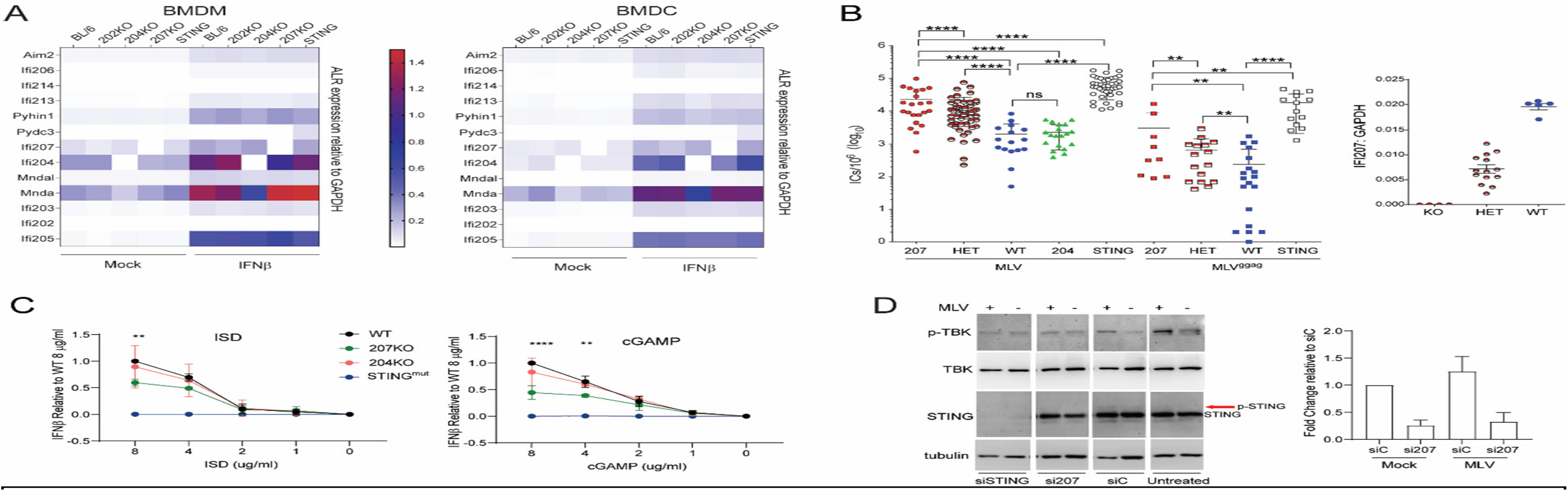
IFI207 is an anti-viral restriction factor. (**A**) *Ifi207* expression in BMDMs and BMDCs. BMDMs and BMDCs from the indicated mice were treated with or without mouse IFN-β for 4h. mRNA expression was quantified by qRT-PCR. Means from 2 independent experiments performed with duplicate or triplicate samples per treatment were plotted. (**B**) Pups of the indicated genotype were infected with 2000 pfu of MLV (MMLV or MLV^gGag^). At 16 dpi, splenic viral titers were determined on NIH3T3 cells. 207 KO, WT and HET pups were derived from IFI207 heterozygote crosses. Right panel: RT-qPCR analysis to verify Ifi207 expression levels in mice. Symbols represent individual mice. Significance was determined by two-tail Mann-Whitney test (**, *P*<0.003; ****, *P*<0.0001). (**C**) BMDMs from the indicated mice were transfected with the indicated concentrations of cGAMP or ISD for 4hr. mRNA expression was quantified by qRT-PCR. Shown are the averages of 3 independent experiments. Two-way ANOVA was used to determine significance (****, *P*<0.0001). (**D**) NIH3T3 cells were transfected with siRNAs targeting IFI207 or STING. Non-targeting (siC) siRNA was also used. At 48h post transfection, cells were infected with MLV^gGag^ for 2h (MOI=5). Protein expression was examined by western blot with the indicated antibodies. Right panel: Ifi207 KD was verified by RT-qPCR. Average +SD from 2 independent experiments is shown.

### IFI207 is an antiviral restriction factor in mice

Since the *Ifi207* KO was gene-specific and did not affect the expression of other *Alr* members, we tested its role in vivo. We infected newborn IFI207 and IFI204 KO mice, as well as mice heterozygous for the Ifi207 KO allele. STING^mut^ mice, which we showed previously are highly susceptible to MLV infection, were infected for comparison (14, 17, 25). Splenic viral titers were measured at 16 dpi. We found that titers were significantly higher in both IFI207 KO and heterozygous mice compared to WT mice (Fig. 2B). Similar results were seen in mice infected mice with MLV^gGag^, a virus mutant with an unstable capsid that induces higher levels of IFN than does WT virus; overall titers were lower than in wild type MLV-infected mice, which we showed previously was due to the increased susceptibility of MLV^gGag^ to nucleic acid sensors and APOBEC3-mediated restriction (14). STING^mut^ mice were more highly infected with both wild type MLV and MLV^gGag^ compared to IFI207 KOs, like our previous studies with knockout of cGAS and DDX41, upstream effectors of STING (14, 17). IFI207 heterozygous mice, which express half as much *Ifi207* RNA compared to WT mice, also showed higher levels of infection than WT mice, suggesting haploinsufficiency (Fig. 2B). Finally, although it has been suggested that IFI204 is an anti-retroviral restriction factor, IFI204 KO mice had titers equivalent to WT mice (13). These data demonstrate that IFI207 is one of the factors that controls MLV infection in vivo.

To further determine if the antiviral activity of IFI207 is mediated by the STING pathway, we next tested whether IFI207 altered STING-mediated type 1 interferon induction. BMDCs from BL/6, IFI207 KO, IFI204 KO and STING^mut^ mice were transfected with increasing amounts of interferon stimulatory DNA (ISD) and or cGAMP, the activating ligand of the STING pathway. Three hours after transfection, IFN-β RNA levels were measured. The IFN response to ISD and cGAMP was significantly reduced in Ifi207 KO BMDMs, while the response by BL/6 and IFI204 cells was the same (Fig. 2C). STING^mut^ cells had no response (Fig. 2C).

The decreased response to STING ligands in IFI207 KO cells was also seen at the protein level. BMDCs from IFI207 KO and WT mice were transfected with ISD, and STING, phospho-STING, phospho-TBK1, and TBK1 were examined by western blot. The levels of all STING, phospho-STING and phospho-TBK1 were reduced in the IFI207 KO cells compared to WT cells (Fig. S5).

We also used siRNA knockdown to deplete IFI207 in NIH3T3 cells and infected them with MLV^gGag^. Knockdown of Ifi207 decreased STING expression and caused lower levels of phospho-STING and phospho-TBK (Fig. 2D). Knockdown of STING also resulted in loss of phospho-TBK in response to infection.

### IFI207 is both nuclear and cytoplasmic

To better understand the function of the S/T repeats, we cloned the *Ifi207* coding regions from RNA isolated from the spleens of C57BL/6N, AIM2 KO and DBA/2J mice as representatives of the variants with different numbers of S/T repeats (Ifi207_BL/6_, Ifi207_DBA_ and Ifi207_129_; Fig. 1C and see below). AIM2 KO mice, whose *Alr* locus is identical to 129P2 mice, were used because 129P2/OlaHsD mice are no longer available8. In addition, for some experiments we used molecular clones of *Ifi203* and *Ifi204* derived from C57BL/6J mice for comparison (6, 14).

Recently it has been suggested that IFI16 restricts HIV-1 infection by interfering with the transcription factor Sp1, thereby suppressing transcription of the integrated provirus (13). This function required nuclear localization of IFI16. IFI204 similarly reduced HIV virus production. To determine where IFI207 might be located within the cell, we first performed fractionations of cells transfected with IFI207_BL/6_, IFI207_DBA_, IFI207_129_ expression vectors in the presence and absence of STING expression plasmids. While a large fraction of IFI207 was in the nucleus, there were also significant amounts in the cytoplasm, independent of the number of repeats; the BL/6, 129 and DBA constructs all showed both nuclear and cytoplasmic localization (Fig. 3A). However, STING also appeared in the nuclear fraction, likely due to its association with perinuclear membrane fractions (28,29). Some of the IFI207 found in the nucleus may be due to a similar association (see below).

**Fig. 3.**
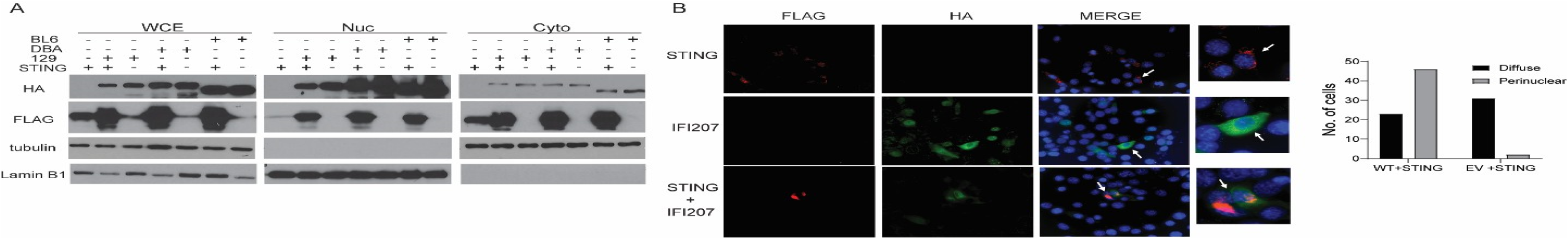
Subcellular localization of IFI207. (**A**) HEK293T cells were transfected with full length IFI207 expression constructs with or without STING expression constructs. Cell lysates were fractionated and proteins were analyzed by western blotting with the indicated antibodies. WCE, whole cell extract; Cyto, cytoplasm; Nuc, nucleus. (**B**) NIH3T3 cells were transfected with full-length IFI207 expression constructs with or without STING expression constructs. Cells were fixed and stained with antibodies directed against expression construct tags. Images were acquired by fluorescence microscopy. Image brightness was manually adjusted to better visualize STING in the STING-alone panels because of the lower levels of STING. White arrows indicate cells that were selected to show enhanced details by zoom. Shown to the right is quantification of STING staining in the presence or absence of IFI207 (diffuse or highly localized/perinuclear) (see Materials and Methods for details).

Comparable results were obtained by immunohistochemistry (IHC); IFI207 was detected in both the nucleus and cytoplasm in the presence and absence of STING (Fig. 3B). STING immunostaining was brighter in the presence of IFI207, confirming the higher levels. Interestingly, STING immunostaining in the absence of IFI207 was more diffuse than in its presence, where it appeared highly concentrated near the nucleus (Fig. 3B).

### Co-expression of IFI207 and STING stabilizes STING

We showed previously that both IFI203 and IFI205 bound STING and that this had functional consequences for activation of the STING pathway in response to exogenous and endogenous retroviruses/retroelements, respectively (8, 14). To determine if IFI207 also bound to STING, we co-transfected flag-tagged STING and all 3 full-length expression constructs (IFI207_BL/6_, IFI207_DBA_ and IFI207_129_) as well as an HA-tagged IFI204 vector and used anti-HA antibodies for immunoprecipitation and western blots. All 3 IFI207 constructs co-immunoprecipitated with STING (Fig. 4A). In contrast, IFI204 weakly co-immunoprecipitated with STING, although it was expressed at much higher levels than any of the IFI207 constructs (Fig. 4A).

**Fig. 4.**
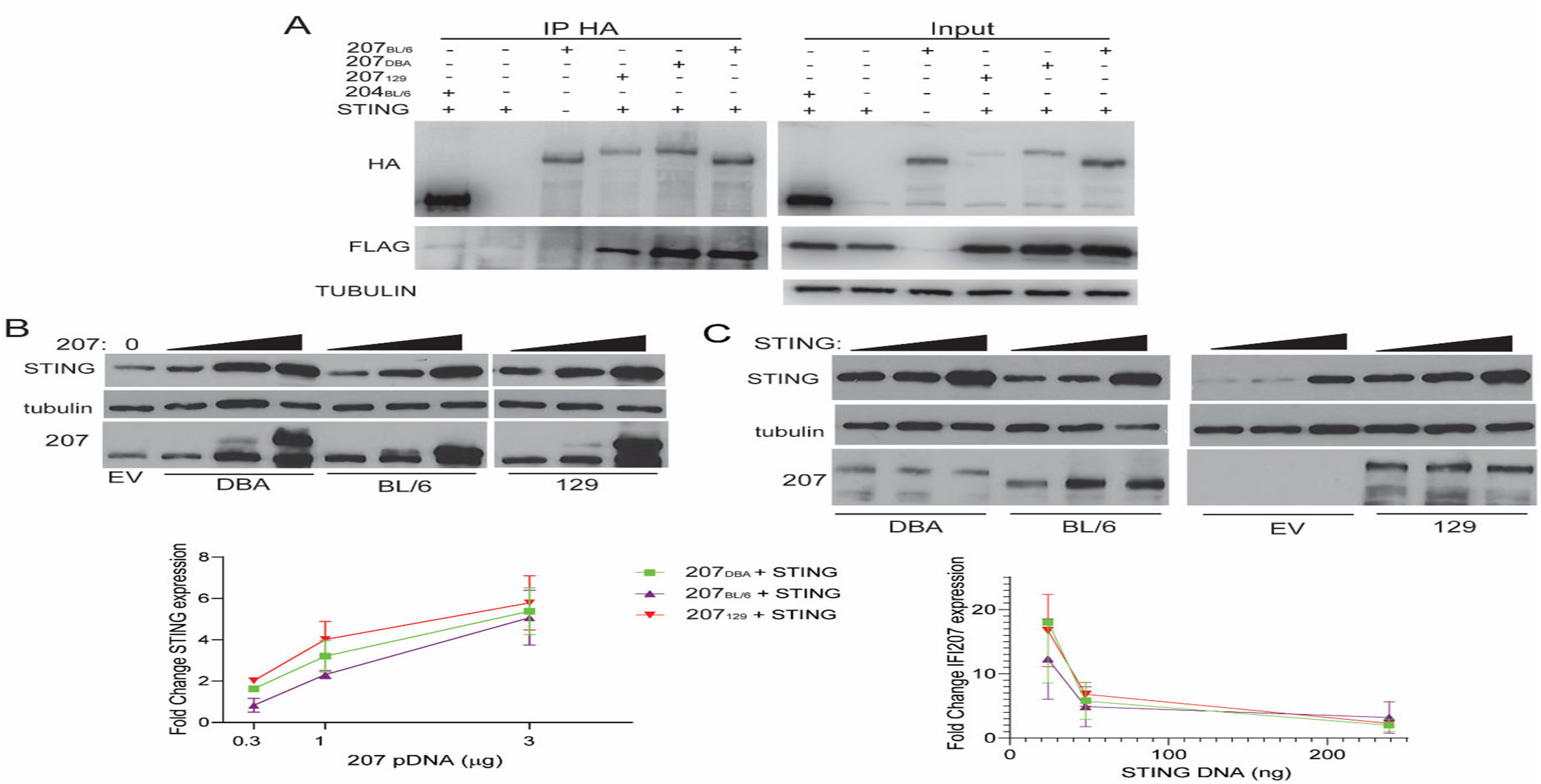
IFI207 stabilizes STING. (**A**) HEK293T cells were transfected with full length HA-tagged IFI207_129_ IFI207_B6_, IFI207_DBA_, IFI204 and flag-tagged STING expression constructs. Anti-HA was used to immunoprecipitated IFI207 in the transfected cell extracts, and western blots with the indicated antibodies were performed. (**B**& **C**) HEK293T cells were transfected with full length IFI207 expression constructs increasing amounts of IFI207 (**B**) or increasing amounts of STING (**C**) expression plasmids. Protein expression was examined by western blot with the indicated antibodies. Experiments were done in triplicate and quantified by densitometry analysis in ImageJ (below). Abbreviations: EV, empty vector.

We observed that even though equal amounts of STING plasmid were transfected in the presence or absence of IFI207, there seemed to be more STING in extracts when IFI207 was present (input, Fig. 4A). This suggested that IFI207 might be stabilizing STING. To determine if this was the case, increasing amounts of the IFI207 expression plasmids were co-transfected with an expression vector containing flag-tagged STING, and the levels of STING were examined by western blot. Co-expression of any IFI207 plasmids stabilized STING protein expression compared to empty vector (Fig. 4B). Moreover, both the IFI207_DBA_ and IFI207_129_ constructs stabilized STING more effectively than the IFI207_BL/6_ construct (Fig. 4B). Increasing amounts of STING plasmid were also co-transfected with constant amounts of the IFI207 plasmids, and again, all 3 IFI207 constructs stabilized STING. Interestingly, while IFI207 stabilized STING, increasing STING had no effect on IFI207 levels (Fig. 4C).

We also examined whether reducing endogenous IFI207 levels altered basal levels of STING in NIH3T3 cells. Transfection of IFI207 siRNAs into these cells reduced basal STING levels compared to control siRNA-transfected or untransfected cells (Fig. S6A).

### The S/T-rich repeat region is responsible for STING stabilization

Although there were minor sequence differences in the PYD and HIN domains of the Ifi207 genes found in BL/6, 129 and DBA mice, the major difference we found was in the sequence of the repeat region (Table S2). To assess whether the repeat region played a role in STING stabilization, we made constructs in the backbone of each allele that lacked all the S/T repeats (R1-IFI207_BL/6_; R1-IFI207_DBA_; R1-IFI207_129_; Fig. 5A). These 3 constructs, as well as the full-length constructs from which they were derived (FL-IFI207_BL/6_; FL-IFI207_DBA_; FL-IFI207_129_), were co-transfected into 293T cells with STING. Expression of all 3 deletion alleles was much higher than full length IFI207 and was comparable between the variants. Nonetheless, deletion of the repeat region abrogated the ability of IFI207 to stabilize STING, since the level of STING after co-transfection with R1-IFI207_BL/6_, R1-IFI207_DBA_ or R1-IFI207_129_ was like that seen with empty vector (EV in Fig. 5A). Moreover, STING dimer formation increased with full-length but not the repeat deletion constructs (Fig. S6B). These data suggest that the repeat region of IFI207 is critical for its ability to stabilize STING.

**Fig. 5.**
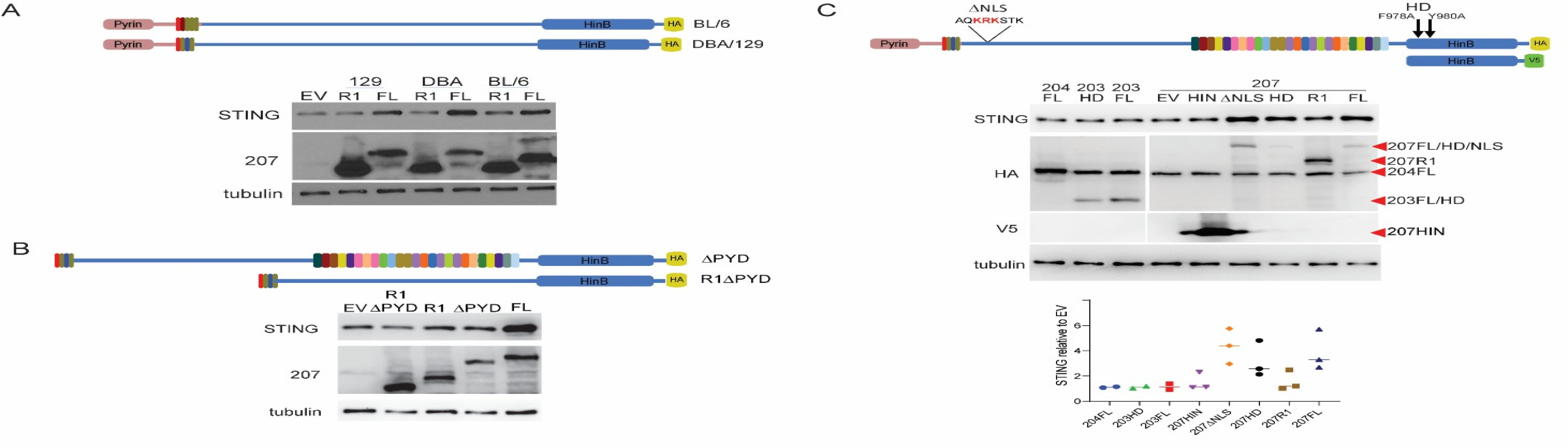
STING stabilization requires the IFI207 repeat region but not DNA binding. (**A**) HEK293T cells were co-transfected with full-length (FL) or repeat deletion (R1) IFI207_129,_ IFI207_B6_, IFI207_DBA_ and STING expression plasmids. Protein expression was examined by western blot with the indicated antibodies. (**B**) The same experiment was carried out with IFI107_DBA_ FL or R1 lacking the PYD domain. (**C**) The IFI207 and IFI203 HD and IFI207 *Δ*NLS mutants are HA-tagged, while the HIN construct contains a V5 tag. HEK293T cells were co-transfected with the indicated IFI207, IFI203, IFI204 and STING expression plasmids. Protein expression was examined by western blot with the indicated antibodies. Bottom panel: triplicate experiments were quantified by densitometry analysis in ImageLab (BioRad). EV=empty vector.

### STING stabilization does not rely on DNA binding or nuclear localization but does require the PYD domain

Previous work showed that the PYD domain of IFI16 was required for its ability to interact with ASC, BRCA1 and STING (30, 31). We thus tested whether IFI207’s PYD domain was also required for STING stabilization. We deleted the PYD domain from both the FL-IFI207_DBA_ and R1-IFI207_DBA_ constructs and co-transfected them with STING. Only the full-length construct, but not the ΔPYD constructs stabilized STING (Fig. 5B). This suggests that although the repeat region was important for STING stabilization, the PYD domain was needed for STING binding, as has been shown for other ALR proteins (30, 31).

We next examined three additional mutants for their ability to stabilize STING: ΔNLS, which deletes a putative nuclear localization signal downstream of the PYD; HD, with alanine substitutions at amino acids 978 and 980 in FL-IFI207_DBA_, which we showed previously abrogated nucleic acid binding when introduced into IFI203 (14); and HIN, which encodes only the HIN domain (Fig. 5C). The ΔNLS and HD constructs showed decreased nuclear localization (Fig. S7). We also subcloned the PYD domain but saw no stable expression of protein (not shown). We co-transfected expression constructs encoding these mutants as well as FL-IFI207_DBA_ and the R1 mutant with STING and examined STING stability by western blots. Constructs bearing IFI204, IFI203 and the DNA binding mutant IFI203 HD were also co-transfected with STING. We found that both IFI207 ΔNLS and HD mutants but not the HIN domain construct stabilized STING (Fig. 5C). Neither the IFI204 nor the IFI203 constructs stabilized STING.

To determine whether any of these regions also contributed to STING binding, we also carried out co-immunoprecipitations on extracts from cells co-transfected with STING and IFI207_DBA_ or the mutants on the DBA backbone. Although they were expressed at much lower levels than the R1 or HIN proteins, the FL, ΔNLS and HD mutants but not the R1 or HIN domain constructs efficiently bound STING (Fig. 6A). As we showed previously, IFI203 also bound to STING, albeit less effectively than repeat-containing IFI207 and at levels similar to the R1 mutant (Fig. 6A) (14).

**Fig. 6.**
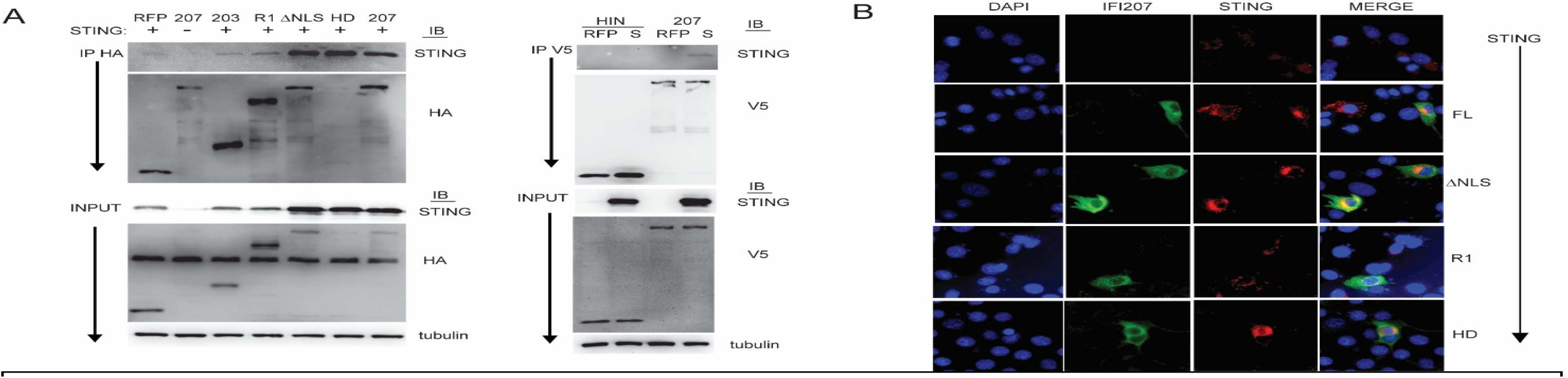
DNA binding is not required for STING stabilization. (**A**) Mutant or WT IFI207_DBA_ or IFI203 expression plasmids were co-transfected with STING in 293T cells and co-immunoprecipitated. Protein expression was analyzed by western blot with the indicated antibodies. (**B**) NIH3T3 cells were transfected with full length or mutant IFI207 expression constructs and STING expression constructs. Cells were fixed and stained with antibodies directed against the expression construct tags. Images of cells transfected with IFI207 constructs without STING are in Fig. S7 and image quantification in Fig. S9.

We also tested whether binding of reverse transcripts correlated with IFI207’s ability to stabilize STING. We showed previously that DDX41 and IFI203 but not the DNA binding mutant, IFI203 HD, precipitated MLV reverse transcription products from infected cells (14). 293T MCAT cells that stably express the MLV entry receptor ATCR1 were transfected with IFI207_DBA_-FL, -R1 and -HD, as well as IFI204, IFI203, and DDX41 and then infected with MLV^gGag^, which, because the capsid dissociates more rapidly than wild type virus, produces high levels of early reverse transcripts (32). At 4 hours post-infection, the ability of the different constructs to precipitate MLV reverse transcripts was determined. IFI207_DBA_-FL and -R1 strongly bound MLV reverse transcripts (strong stop region), while IFI207 HD weakly bound viral DNA (Fig. S8). IFI204 and IFI203 also weakly bound viral DNA, as we previously showed (14). These data show that nucleic acid binding/activation is not required for IFI207 stabilization of STING, at least under conditions of over-expression.

Next, we studied the co-localization of the mutants by immunofluorescence. Both the ΔNLS and HD mutants were more frequently localized in the cytoplasm, while like the WT protein, the R1 mutant was seen in both the nucleus and cytoplasm (Fig. 6B and Fig. S7 and S9A). The WT and ΔNLS but not the HD or R1 constructs caused STING to be found in concentrated perinuclear foci (Fig. 6B and Fig. S9B).

## DISCUSSION

Pathogens infect different cell types and tissues and many replicate in distinct cellular compartments. TLRs, ALRs, cGAS, and other sensors mediate innate immune responses to pathogens. The need for targeted responses to different pathogens has likely led to this expansion of receptors with different signaling pathways. These specific sensing and signaling pathways are important targets for pathogens, and as a result, greatly accelerate host-pathogen co-evolution. Indeed, in house mouse strains, ALRs, IFNs and even IFN receptors are observed as “strain-specific diverse regions” (SSDRs) with high sequence diversity (32). Some of the SSDRs are among the most dynamic regions in the genome. Both genome recombination (33), and non-crossover gene conversions (34) between large gene families with two divergent haplotype genomes will create novel gene structures. Unlike housekeeping genes, newly created immune-related genes have a higher chance of being beneficial because of the “red queen hypothesis” and drifting in the population (34). On the other hand, the destruction of an existing gene by recombination or gene conversion is more likely to be tolerated when it is part of a gene family because other family members may have (semi-) redundant functions. As a result, SSDRs are possibly also hot spots of new gene generation.

The *Alr* locus is a typical SSDR. It is found in most mammalian genomes, except for some bat species (35). The ALRs belong to one family of innate immune sensors that may have diversified in response to the selective pressures of pathogens (6-9, 36). We show here that several new *Alr* family members have recently been introduced into the genomes of the *Mus* genus, probably by one or a series of genome recombination events. Novel ALR members like MNDAL and IFI207 have acquired new roles in variable biology processes: MNDAL has become a cell growth regulator (37), and as we show here, IFI207 has acquired a key role in the STING mediated antiviral signaling pathway, using non-redundant and different mechanisms than the other ALRs. These new members may have modified the functional capacity of mouse ALRs, because in rats and humans, fewer ALR members are sufficient for normal immune function. For example, humans have *AIM2*, *IFI16*, *PYHIN1*, and *MNDA*, while rats have *Rn7* (*AIM2*), *Rhin2* (*Ifi206*), *Rhin3* (*Ifi214*), *Rhin4* (*Ifi203*) and *Rhin5* (*Mnda*) (Fig. 1B). Thus, no *Ifi207* orthologues are required in these species.

Although the entire *Alr* locus shows great diversity among different house mouse strains, *Ifi207* was strikingly diverse. We found that this gene contains a large PYD-proximal repeat region that varies extensively among inbred and wild mice strains. We analyzed the repeat region sequence in multiple mouse strains and found a large collection of different sequences of repeating units in the IFI207 repeat region. In addition to sequence variation of the repeat region, these mice also had different numbers of repeating units. Apart from this repeat region, the rest of the protein sequence was almost identical in different inbred strains, suggesting that the repeat region specifically has been a target of selective pressures. The combination of repeat numbers and sequences in *Ifi207* is potentially unlimited. A better understanding of how the Ifi207 repeat region has evolved might offer clues about its role as an innate immune effector as wild mice encounter a wide array of pathogens that may have shaped its evolution.

The repeat region, which is predicted to form a coiled structure similar to a leucine zipper, makes IFI207 unique among the ALRs. The length of the repeat region also affected binding to STING and its stabilization. The IFI207 proteins encoded in DBA/2 and 129 mice, which have 25 and 24 repeats, respectively, stabilized STING more effectively than did the BL/6 protein, which has only 12 repeats. This suggests that these repeats expanded and were subsequently retained in the encoded protein, likely because of exposure to pathogens by the progenitor mice from which different inbred strains were derived. Several ALRs, including IFI16, IFI203, IFI204 and IFI205 have been shown to bind STING, but these proteins lack the extended repeat region, and as we show here, bind STING to the same extent as IFI207 with no repeats (8, 14). Thus, although the PYD domain of different ALRs may confer binding to STING, it is the IFI207 repeat region that confers its stability and the enhanced response to different ligands and viruses that act in the STING IFN response pathway.

A previous study suggested that knockout of the entire Alr locus in mouse macrophages had no effect on the response to ISD (Gray et al., 2016). However, a more recent study showed that ALR KO mice were more highly infected with MLV (Hotter et al., 2019). A number of studies have implicated IFI204 in the response to herpesviruses, polyomavirus and several bacteria such as *Francisella* and *Staphyloccocus*, and IFI16 is important in human cells for the innate immune response to *Listeria*, herpesviruses and HIV (10, 13, 16, 38-41). We showed previously that IFI203, in conjunction with DDX41, recognized MLV and that DDX41 KO mice were as susceptible to MLV infection as cGAS KO mice (14). We show here that IFI207 alters the response to DNA, cGAMP and MLV. Since MLV has been in *Mus* species for at least 1 million years, it is not surprising that there is an arsenal of restriction factors in mice that control MLV infection, including multiple ALRS (42). Whether IFI207 KO mice are more susceptible to infection with other pathogens that activate the STING pathway is currently under investigation.

IFI205 and IFI16 act as transcriptional repressors of Asc and integrated HIV proviruses, respectively, and this activity requires nuclear localization (11, 13). Although IFI204 was shown to act as an anti-MLV and -HIV restriction factor by suppressing proviral transcription, we did not see any effect in IFI204 KO mice on infection levels, suggesting that it is not anti-viral in vivo. IFI207 also has an NLS, but we show that its ability to stabilize STING was independent of this sequence. Interestingly, at least in over-expression studies, the mutant lacking repeats still bound bind MLV reverse transcripts but failed to stabilize STING, while the HD mutant that has minimal nucleic acid binding still showed STING-stabilizing activity. The role of the nucleic acid binding domain in IFI207 function thus remains to be determined.

There is incredible diversity in the *ALR* locus among mammalian species. For example, the horse locus has 6 ALR genes, dogs and pigs have 2, cows have 1, rats have 5, humans have 4 genes, and bats have only a truncated AIM2 gene, all of which are encoded in a single locus which sits between the cell adhesion molecule 3 (CADM3) and spectrin alpha, erythrocytic 1 (SPTA1) genes (6-9, 35). In addition to their roles in innate immunity, several ALRs have been implicated in adipogenesis (43, 44), autoimmune diseases like lupus erythematosus (45, 46) and inflammatory responses (11, 47). Identification of the pressures that shaped the locus in different species is thus important for understanding the many biological processes in which these genes play a role.

## MATERIALS AND METHODS

### Mice

Mice were bred at the University of Illinois at Chicago (UIC). C57BL/6N, DBA2/J, BALB/cJ, STING^mut^ (C57BL/6J-Sting1gt/J), AIM2 KO (B6.129P2-Aim2Gt(CSG445)Byg/J) and CMV-Cre (B6.C-Tg(CMV-cre)1Cgn/J) mice were originally purchased from The Jackson Laboratory. IFI207 KO mice (C57BL/6N-IFI207em1(IMPC)Wtsi) were generated by targeted deletion by zygote injection of 4 gRNAs, two flanking each side of exon 3, to induce a frameshift null allele of Ifi207. The embryos used for these experiments were C56BL/6NJ, and after germline transmission, the colony was maintained on this background. Full details of the gRNAs used for gene disruption are provided in Table S3A. IFI202 (Ifi202btm1(KOMP)Mbp/MbpMmucd) and IFI204 KO mice (Ifi204tm1a(KOMP)Wtsi/MbpMmucd) were derived from sperm purchased from the Mouse Biology Program and UC Davis. The IFI204 mice were initially crossed with CMV Cre mice to generate knockout strains and then backcrossed to remove the CMV Cre transgene (Fig. S4). Genotyping primers for the Ifi207, Ifi202 and Ifi204 knockout alleles are in Table S3B. cGAS KO mice (Mb21d1tm1d(EUCOMM)Hmgu) were originally provided by Michael Diamond and Skip Virgin (48). 129P2/OlaHsd DNA was isolated from mice originally purchased from Harlan (8). M. m. castaneus mice were purchased from the Jackson Laboratory. All mice were housed according to the policies of the Animal Care Committee (ACC) of the University of Illinois at Chicago (UIC); all studies were performed in accordance with the recommendations in the Guide for the Care and Use of Laboratory Animals of the National Institutes of Health. The experiments performed with mice in this study were approved by the UIC ACC (protocol no. 21-125).

### Sequence analysis

The assemblies of non-mouse species are downloaded from NCBI (https://www.ncbi.nlm.nih.gov), with access names listed in Table S1 and S2. Whole genome de novo assembly of 13 mouse strains, Mus spretus and Mus caroli were provided by the Phase 3 mouse genomes project (MGP) from EMBL-EBI (unpublished data, available on ENA database with access number PRJEB47108). The assemblies were produced using third generation long reads (PacBio Sequel / PacBio Hifi, BioNano®) and Hi-C. For the additional mouse strains, the alleles of Ifi207 were assembled from whole genome Illumina data or the transcriptomes of IFN-α/γ induced diaphragm fibroblast cells. Samples were provided by Institute of Vertebrate Biology CAS. For a further confirmation, the sequences of HIN-side repeat regions were double confirmed by Sanger sequencing of genome amplicons, with the primers described in Table S6. Full-length sequences of DBA and 129 IFI207 were obtained from the cloned cDNAs (see Plasmids and Transfection). Genbank acquisition numbers for the re-sequenced repeat regions and full-length clones are listed in Supplementary Table S4. For more details, please see Plasmids and Transfection.

### Cells

NIH3T3 and HEK293T cells were cultured in Dulbecco’s Modified Eagle Medium (DMEM) supplemented with 10% FBS, 2mM L-glutamine (Gibco), 100 U/ml penicillin (Gibco), and 100 mg/ml streptomycin (Gibco). Cell cultures were maintained at 37°C with 5% CO2. MCAT cells (293T cells expressing ATCR1, the MLV entry receptor50) were cultured in DMEM supplemented with 8% donor bovine serum, L-glutamine, penicillin/ streptomycin and G418 (10 mg/ml) (Goldbio).

### BMDM and BMDC cultures

Bone marrow was harvested from hind limbs of mice. To differentiate BMDMs, cells were cultured for 6-7 days on Petri dishes in DMEM containing 10% FBS, 2 mM L-glutamine, 100 U/ml penicillin, 100 g/ml streptomycin, 0.1 mM sodium pyruvate, 25mM HEPES (Gibco), and 10 ng/ml murine M-CSF (Gibco, PMC2044). Fresh media containing 10 ng/ml M-CSF was added on day 3. On day 6 or 7, BMDMs were collected for experiments. To differentiate BMDCs, bone marrow cells were treated with ACK, and 2×106 cells were seeded in the center of Petri dishes in 10ml RPMI containing 10% FBS, 2 mM L-glutamine, 100 U/ml penicillin, 100 ⍰g/ml streptomycin and 50 ⍰M β-mercaptoethanol plus 20 ng/ml murine granulocyte-macrophage colony-stimulating factor (GM-CSF) (Peprotech, 315-03). On day 4, 10 ml fresh media containing 20 ng/ml GM-CSF was added to plates. On day 8, half of the culture volume was carefully removed to avoid disrupting the cells clustered in the centers of dishes, and 10 ml fresh media with 20 ng/ml GM-CSF was added to plates. Non-adherent cells clustered in centers of wells were collected by gentle pipetting for experiments on day 10. At time of collection, BMDMs and BMDCs were stained for F4/80, CD11b, or CD11c and analyzed by flow cytometry (LSR-Fortessa flow cytometer; BD Biosciences). Macrophage and DC cell populations were >80% pure.

### Expression Analysis

BMDMs and BMDCs from BL/6, STINGmut, IFI202, IFI204 and IFI207 KO mice were treated with or without 100 U mouse IFN-β (PBL Assay Science, 12405-1). Four hours after treatment, RNA was harvested, and Alr gene expression was analyzed by qRT-PCR. RNA was isolated by RNeasy Mini Kit (QIAGEN 74106) with on-column DNase digestion using RNase-Free DNase Set (QIAGEN 79254). cDNA was synthesized with random hexamers from the Superscript III First-Strand Synthesis System (Invitrogen 18080051). qPCR was performed using Power SYBR Green PCR Master Mix (Applied Biosystems 4367659) with appropriate primer sets on a QuantStudio 5 Real-Time PCR System (Applied Biosystems). Primers for qRT-PCR analysis of Alrs, Ifnb, actin, and gapdh have been previously described (Table S5) (8, 14). MLV strong-stop primers for have been previously described (49).

### Stimulation of BMDMs and BMDCs

BMDMs were transfected with 8, 4, 2, 1, or 0 μg/ml interferon stimulatory DNA (ISD) (InvivoGen) or 2’3’-cGAMP (Sigma) for 3h. RNA was harvested, and mRNA expression was analyzed by qRT-PCR. Transfections were done using Lipofectamine 3000 (Invitrogen). For western blot analysis, BMDCs were transfected with ISD for 2 h. After 2 h, culture media was removed and replaced with fresh media, and samples were incubated for an additional 4, 8, or 12 hours. At indicated time points, BMDCs were collected in PBS, pelleted, lysed in radioimmunoprecipitation (RIPA) buffer (50mM Tris pH 8, 150mM NaCl, 1mM EDTA, 1% Triton X-100, 1% sodium deoxycholate, and 0.1% SDS), and prepared as described in the western blot section below.

### Virus

Wild-type Moloney MLV and MLV^gGag^ were harvested from supernatants of stably infected NIH3T3 cells. Supernatants were centrifuged for 10mins (3000rpm) to remove cellular debris, and then filtered through a 0.45 μm filter (Millex Millipore) and treated for 30 min at 37°C with 20 U/ml DNase I (Roche). Virus was pelleted through a 25% sucrose cushion, resuspended in supplemented 2% FBS DMEM, snap frozen on dry ice with ethanol, and stored at −80°C. Virus titers were determined by an infectious center (IC) assay on NIH3T3s. One day before titering, 8×104 cells were plated in 6-well tissue culture plates and incubated overnight at 37°C. Ten-fold serial dilutions of virus stocks were prepared with polybrene (8 μg/ml) in supplemented DMEM and added to cells. Infected cells were incubated at 37°C for 2 h with rocking every 15 min. After 2 h, viral dilutions were aspirated, and 2.5 ml media was added to each well. Plates were incubated for 3 days at 37°C. After 3 days, plates were washed in PBS and stained with monoclonal 538 Env antibody (1:300) for 1 hr at 4°C with rocking. The plates were washed with 2% FBS in PBS and stained with Goat anti-mouse IgG/IgM Alexa Fluor 488 conjugated secondary antibody (Invitrogen A-10680) diluted in 2% FBS in PBS (1:300) for 45 mins at 4°C with rocking. Plates were washed and infectious centers were counted by microscopy (Keyence BZ-X710 and BZ-X analyzer) to determine viral titers. For some stocks, viral mRNA was quantified by RT-qPCR with primers for env and viral proteins were confirmed by western blotting with MLV antisera, as previously described (50).

### In vivo infections

IFI207 heterozygous mice were crossed, and their newborn litters consisting of knockouts and heterozygotes, as well as WT mice, were used for infection. Genotyping was done after virus loads were measured. Additional BL/6 pups were also infected and included in the WT group. STING^mut^ and IFI204 KO mice were generated by homozygote matings. Mice were intraperitoneally injected with 2×10^3^ PFU MLV at 0-48hrs-post birth. At 16 days post-infection, splenocytes were harvested and treated with ACK Buffer for RBC lysis. Tenfold dilutions of the cells were co-cultured with NIH3T3 cells for 3 days to assess viral titers by IC assay.

### Plasmids and Transfections

Expression plasmids for DBA and 129 IFI207 variants were generated by PCR amplification of cDNA synthesized from splenic RNA of DBA2/J and AIM2 KO mice. AIM2 KO were used because 129/P2 mice were no longer available for purchase, and the Alr locus, and therefore Ifi207, in AIM2 KO mice is derived from the parental 129P2/OlaHsd embryonic stem cells in which the knockouts were made (8). An HA tag was added to the C-termini immediately before the stop codon of both IFI207 variants. The PCR products were cloned into pcDNA3.1/Myc-His(+) (Invitrogen). Full-length sequences of DBA and 129 IFI207 were obtained from the cloned cDNAs. BL/6-derived IFI207-HA, IFI204-HA and STING-FLAG plasmids were a gift from Dan Stetson. The IFI203 and IFI203HD plasmids were previously described (14). V5 tags were introduced to all 3 IFI207 variants by PCR amplification of the IFI207-HA expression constructs with primers designed to replace the HA tags with V5 tags. The V5-tagged PCR products were cloned into pcDNA 3.1/Myc-His(+). The myc-tagged DDX41 construct was previously described14. IFI207 R1 mutants, *Δ*NLS, PYD and HD mutants were generated from full length IFI207-HA plasmids and the R1-PYD mutant from R1-IFI207 using the Q5 Site-Directed Mutagenesis Kit (New England Biolabs). IFI207-HIN plasmids were generated by PCR amplification of the HIN domain from full length IFI207-V5 expression plasmids. All cloning primers are described in Table S6.

### Transfections for Protein Expression Analysis

HEK293T cells were transiently transfected using Lipofectamine 3000 for 24 h. For fractionation assays, HEK293T cells were transfected with expression plasmids, collected in ice cold PBS, and fractionated into nuclear, cytoplasmic, and whole cell extracts by the Rapid, Efficient And Practical (REAP) method (51). The REAP method was modified to lyse nuclear extracts in RIPA buffer and to treat nuclear lysates with 3U Micrococcal Nuclease (Thermo Scientific) per μl lysate for 15 min at RT. For IFI207 and STING titration assays, HEK293T cells were transfected with either increasing amounts of IFI207-HA and a constant amount of STING-FLAG or with increasing amounts of STING-FLAG and a constant amount of IFI207-HA. When different amounts of plasmids were transfected within experiments, empty vector (EV) plasmid (pcDNA 3.1/Myc-His(+)) was added to keep the total amount of transfected DNA equal for all samples.

### Immunoprecipitations

For immunoprecipitation (IP) assays, HEK293T cells were transfected with expression plasmids for 24 h using Lipofectamine 3000. Cells were harvested in PBS, pelleted, lysed in RIPA buffer, sonicated, and quantified by Bradford Assay (Bio-Rad). IP samples were prepared with Protein A/G PLUS-Agarose (Santa Cruz sc-2003) or HA-tag and V5-tag Sepharose bead conjugate (3956S and 67476S, respectively). Antibodies and Agarose or Sepharose beads were incubated with cell lysates at 4°C overnight on a rotating rack. Samples were washed 3-5 times in RIPA buffer and centrifuged 3 min (2500 rpm). Samples were eluted in 4X Laemmli buffer plus 5% β-mercaptoethanol.

### siRNA knockdowns

NIH3T3s were transfected with siRNAs (Silencer Select, Ambion by Life Technologies) targeting Ifi207 (s105479, s105481, s202438) and STING (s91056) using Lipofectamine RNAiMAX (Invitrogen). Non-targeting control siRNA was also included (AM4635). Forty-eight hours post transfection, cells were collected and analyzed by western blot analysis or cells were infected with MLVgGag and harvested at 2 hours post infection. Knockdown efficiency was determined by RT-qPCR with primers specific for targets.

### Western blots

For all protein expression analysis, lysates were sonicated on ice, quantified by Bradford Assay, prepared in 4X Laemmli buffer plus 10% β-mercaptoethanol, and boiled for 5 mins. Protease and Phosphatase inhibitor cocktail (Thermo Scientific 78443) was added to all lysis buffers immediately before use. Proteins from cell lysates were resolved by sodium dodecyl sulfate polyacrylamide gel electrophoresis (SDS-PAGE) and then transferred to polyvinylidene difluoride membranes (Millipore IPVH00010). For native blots, lysates were prepared in 4X loading buffer without SDS, and gels were run with Tris-glycine running buffer without SDS. Proteins were detected with primary antibodies against HA-tag (Cell Signaling Technology (CST) C29F4 or Abcam 9110), FLAG-tag (Sigma F1804 or CST 2368), V5-tag (Invitrogen 46-0705), STING (CST D2P2F or D1V5L), lamin B1 (CST D4Q4Z), IκB⍺ (CST 44D4), ⍺-tubulin (Sigma T6199), TBK1 (CST 3013), and phospho-TBK1 (Ser172) (CST D52C2). HRP-conjugated antibodies (CST anti-rabbit or anti-mouse HRP IgG,7074 and 7076, respectively) were used for detection with Pierce ECL Western Blotting Substrate (Thermo Fisher Scientific, 322209) or ECL Prime Western Blotting Reagent (GE Healthcare Amersham, RPN2232).

### Immunohistochemistry

NIH3T3 cells were seeded on glass coverslips in 12-well plates and incubated at 37°C overnight. Cells were transfected with IFI207-HA and STING-FLAG expression plasmids for 24 h with Lipofectamine 3000. To stain, culture media was aspirated, cells were fixed in 4% paraformaldehyde (PFA) for 10 min, washed in PBS for 3×2 min, permeabilized in 0.25% Triton X-100-PBS for 5 min, washed, blocked for 30 min in 10% goat serum/PBS, incubated in primary antibody 2 h, washed, incubated in secondary antibody 2 h, and washed a final time. Coverslips were mounted on slides with VECTASHIELD Antifade Mounting Medium with DAPI (Vector Laboratories, H-1200). Primary and secondary antibodies were diluted in blocking buffer. Primary antibodies used were rabbit anti-HA (Abcam 9110) and mouse anti-FLAG M2 (Sigma, F1804). Goat anti-Rabbit IgG Alexa Flour 568-conjugated (Invitrogen, A-11011) and goat anti-mouse IgG Alexa Fluor 488-conjugated (Invitrogen, A-11001) secondary antibodies were used. Images were captured on a Keyence BZ-X710 at 60X magnification under oil immersion. Image brightness was manually adjusted to better visualize STING. Scoring of images was done in a blinded fashion. Images were scored for IFI207 and STING localization (cytoplasmic, nuclear, or both). The pattern of STING staining was also characterized as concentrated/perinuclear or diffuse. To quantify localization, cells were counted that were IFI207-positive or STING-positive (single transfections) or IFI207-and STING-double positive (double transfections). Cells scored as cytoplasmic, nuclear, or both are represented either as raw cell numbers (Fig. 3B and Fig. S9B) or as the proportion of the total cells counted within each transfection group (Figs. S7 and S9A).

### Nucleic acid pulldowns

Pulldowns were performed as previously described51. Briefly, 293T MCAT cells transfected with pcDNA3.1 (empty vector) or the indicated expression constructs were infected with virus and at 4 hpi, the cells were cross-linked with 1% formaldehyde in media. Cross-linking was quenched with 2.5M glycine, and then extracts were incubated overnight with anti-myc-or anti-HA-agarose beads (Sigma) and G/A-agarose beads (SantaCruz). The beads were washed with high-salt buffer (25mM Tris-HCl, pH 7.8, 500mM NaCl, 1mM EDTA, 0.1% SDS, 1% TritonX-100, 10% glycerol) and with LiCl buffer (25mM Tris-HCl, pH 7.8, 250mM LiCl, 0.5% NP-40, 0.5% Na-deoxycholate, 1mM EDTA, 10% glycerol). The immunoprecipitated nucleic acid was eluted from the beads at 37°C in 100mM Tris-HCl, pH 7.8, 10mM EDTA, 1% SDS for 15 min, and the protein-nucleic acid cross-linking was reversed by overnight incubation at 65°C with 5M NaCl. The eluted nucleic acid was purified using the DNeasy Kit (Qiagen) and analyzed with RT-PCR strong stop primers (SSS) (Table S5) (49).

### Statistical Analysis

All gene expression qRT-PCR analysis experiments were done in duplicate or triplicate with 2-3 technical replicates for each experiment. Each technical replicate was additionally split into triplicates for qPCR analysis. All other experiments were performed at least in triplicate with technical replicates or as indicated in the figure legends. Fluorescence quantifications were done in a blinded fashion. GraphPad Prism was used for statistical analysis. Tests used to determine significance are shown in the figure legends.

## Acknowledgments

We thank David Ryan for help with the mouse breeding and analyzing the immunofluorescence data, and the members of our lab for helpful discussions. Supported by National Institutes of Health Grant R01AI121275 (SRR).

---

**Fig. S1.**
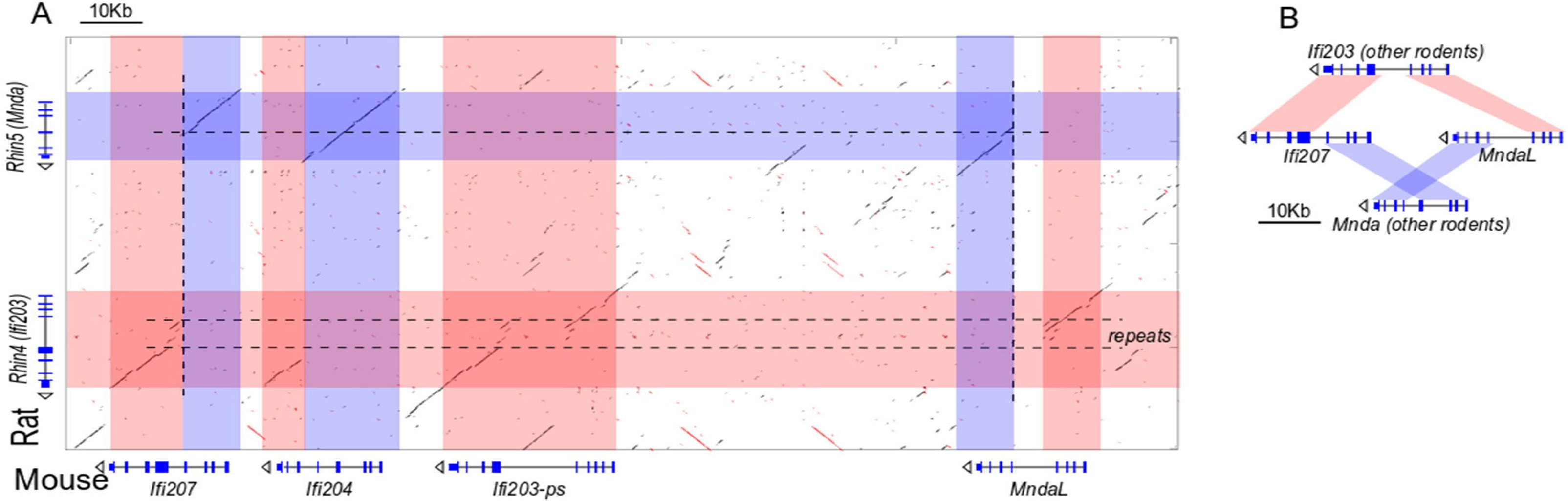
*Ifi207* and *MndaL* in the *Mus* Genus are possibly created by recombination between ancestral *Ifi203* and *Mnda* genes. Using the rat genome, which is similar to most other rodent species, as an example in the dot plot, the structure of mouse *Ifi207*, *Ifi204* and *MndaL* appear to be the result of recombinations between rat *Mnda* (*Rhin5*, indicated with blue shades) and *Ifi203 homologue* (*Rhin4*, red shades). Broken at a repeat rich region, the ancestral *Ifi203* and *Mnda* fused into mice *Ifi207* and *MndaL*.

**Fig. S2.**
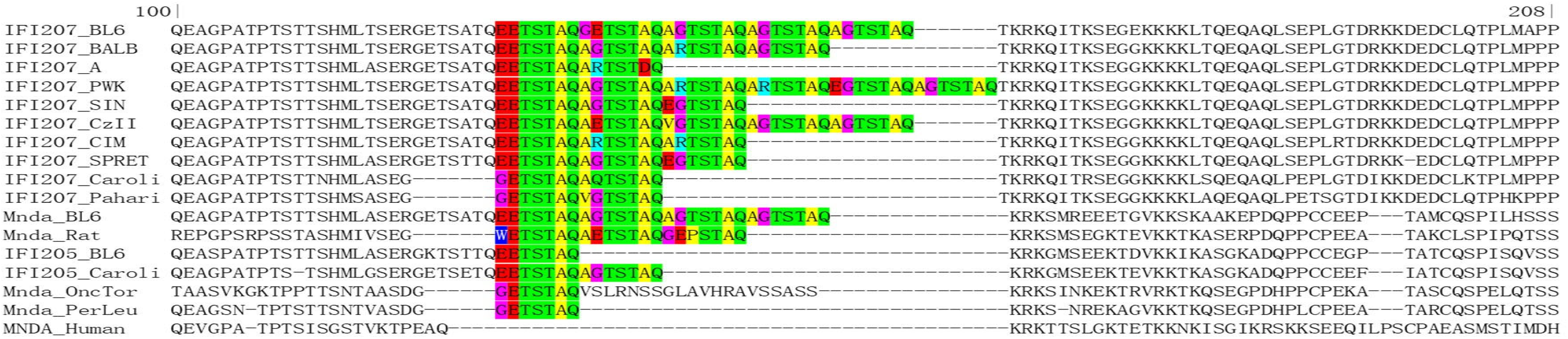
Amino acid sequence around the PYD-side repeat region in human MNDA, rodent IFI207, IFI205 and Mnda. Color codes for the repeat unit: green, polarized (T,S,Q); yellow, non-polarized (A,V,P); cyan, positively-charged (R); red, acidic(E,D); blue (W) and purple (G). Numbers on top indicates residue location in IFI207_C57BL/6J.

**Fig. S3.**
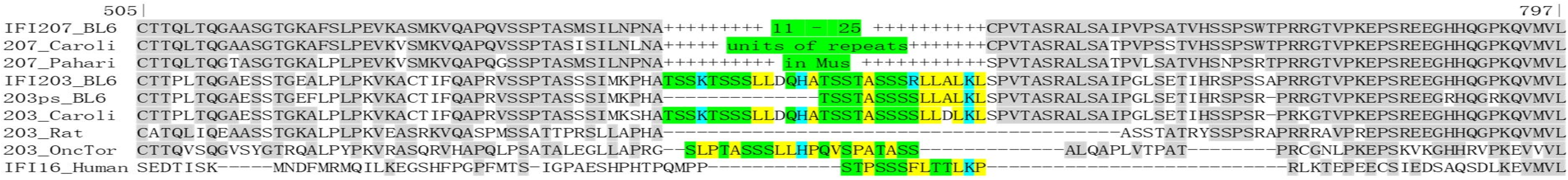
Amino acid sequence around the HIN-side repeat region in human IFI16, rodent IFI203 and IFI207. Rodent IFI203 has 1-2 copies of HIN-side repeat sequence, and human IFI16 also has one copy. Color codes for the repeat unit: green, polarized (T,S,Q); yellow, non-polarized (F,L,A,P,V); cyan, basic (H & K). Numbers on top indicates residue location in IFI207_C57BL/6J.

**Fig. S4.**
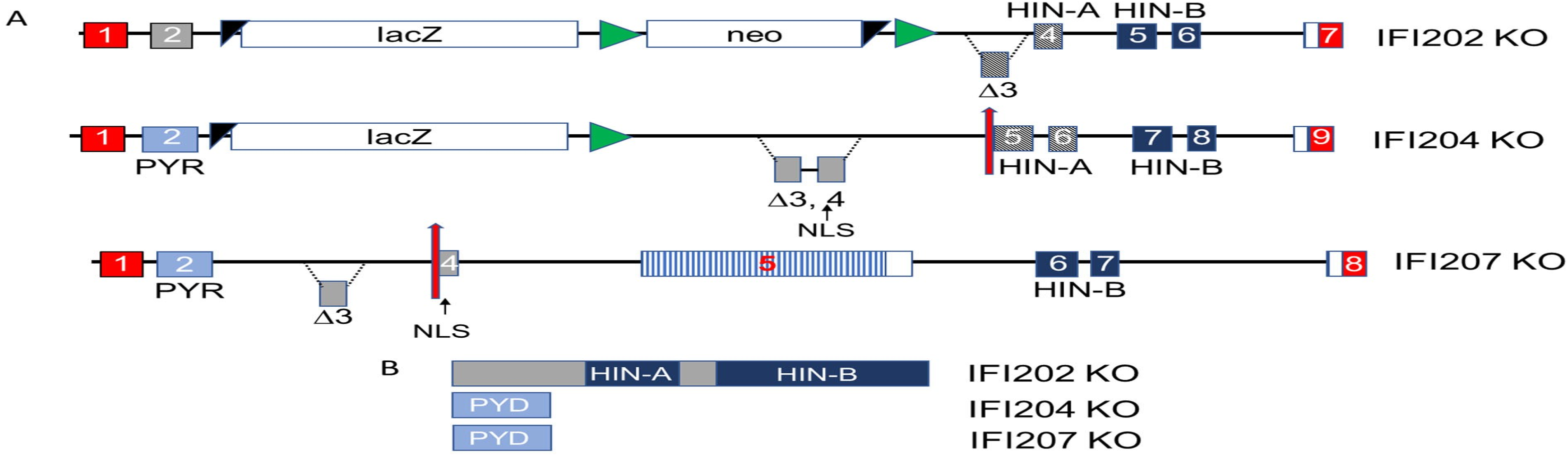
*Ifi202*, *Ifi204* and *Ifi207* KO alleles. (**A**) Diagram of the mouse KO alleles. Not drawn to scale. Exon 3 in the *Ifi202* gene was replaced with a lacZ/neo cassette. This results in a protein lacking exon 3. A similar lacZ/neo cassette with flox sites upstream and downstream of exons 3 and 4 was engineered into the *Ifi204* locus; crossing the parental mice with CMVcre mice resulted in the deletion of these exons and the introduction of a stop codon at the beginning of exon 5 (red arrow). Details of the construction of the parental *Ifi202* and *Ifi204* mouse knockouts can be found on the KOMP Repository website. CRISPR/Cas9 was used to delete exon 3 of the *Ifi207* gene, as described in Methods. This causes a stop codon at the beginning of exon 4. Non-coding sequences are depicted in red. Striped sequences in exon 5 of *Ifi207* represent the repeat region. NLS, nuclear localization signal. (**B**) Predicted proteins from the KO alleles.

**Fig. S5.**
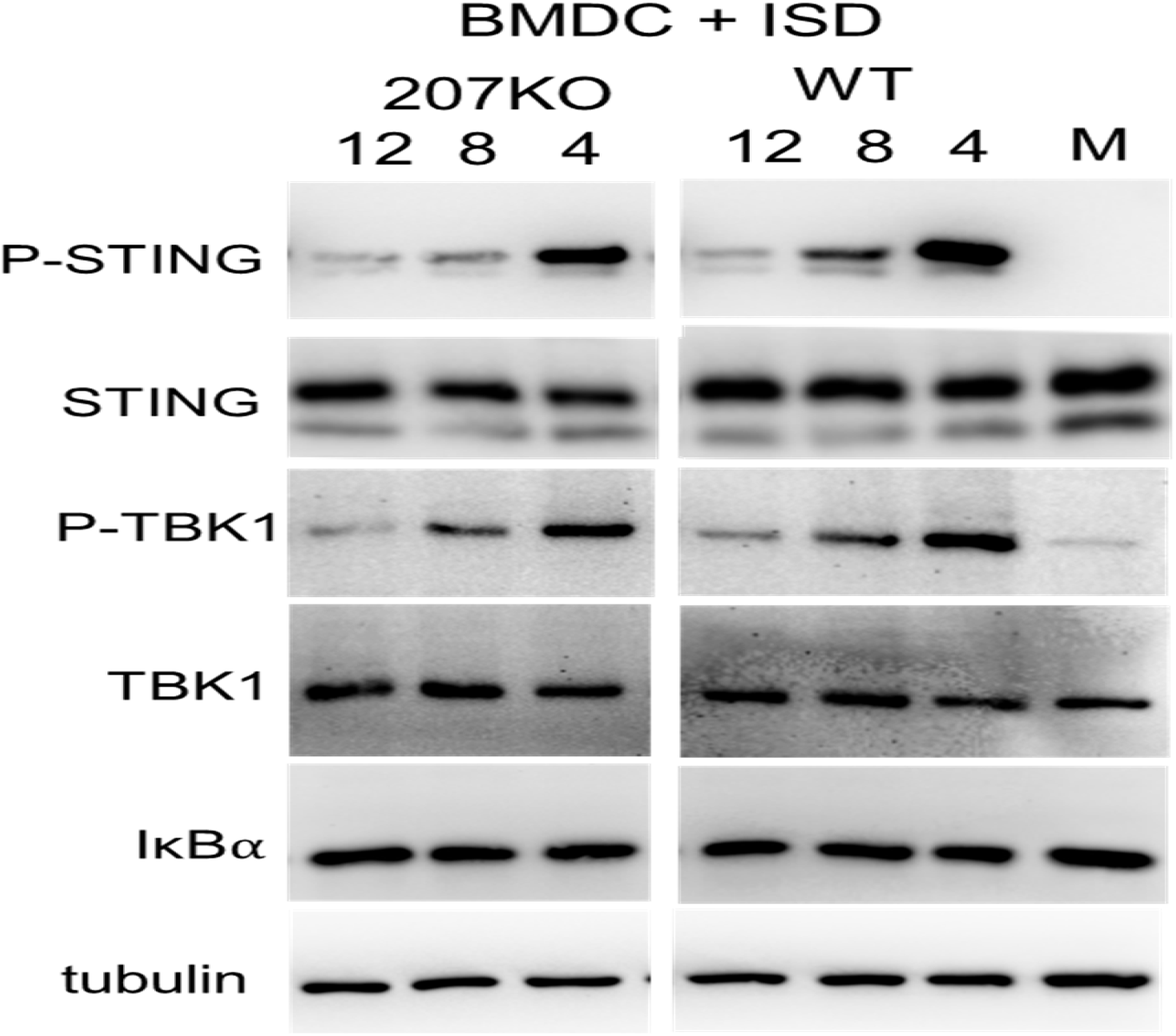
Endogenous STING levels are reduced in Ifi207 KO cells. BMDCs were transfected with ISD 4, 8 and 12 hr and protein expression was analyzed by western blot with the indicated antibodies. Blots were stripped and probed with anti-tubulin to control for loading.

**Fig. S6.**
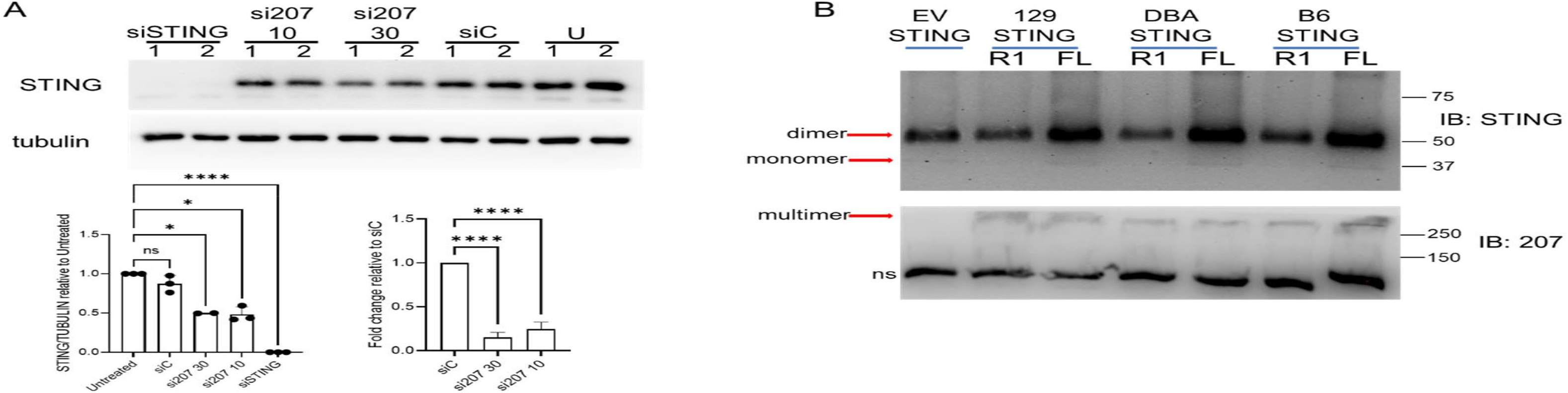
IFI207 stabilizes STING. (**A**) NIH3T3 cells were transfected with the indicated siRNAs, and western blots for endogenous STING were performed. Two concentrations of the 207 siRNA were used (10 and 30 pmole). Duplicate experimental replicates are shown. Bottom: Quantification of western blots from 3 independent experiments (left) and Ifi207 knockdown verification by RT-qPCR (right). Average + SD from 3 independent experiments is shown. One-way ANOVA was used (*, *P* ≤0.04; ****, *P* ≤ 0.0001; ns, not significant). U, untransfected. (**B**) Full-length IFI207 increases STING dimers. 293T cells were co-transfected with IFI207 and STING expression plasmids. Lysates were run on native gels and protein expression was examined by western blot with the indicated antibodies. FL, Full length IFI207; R1, Repeat mutant IFI207; EV, empty vector; ns, non-specific.

**Fig. S7.**
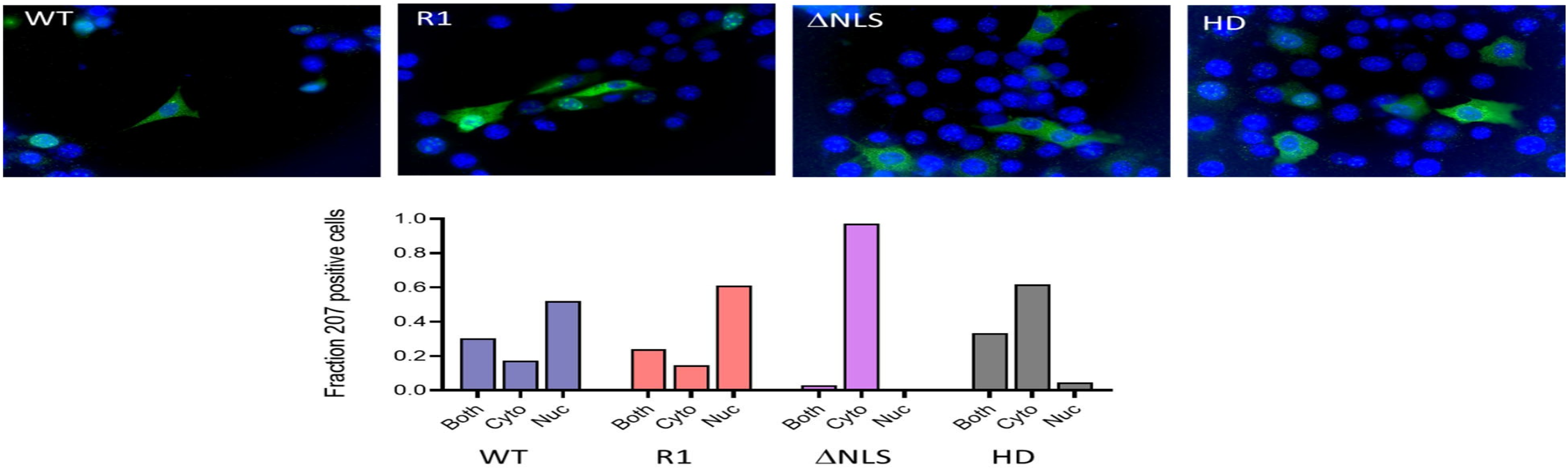
Subcellular localization of IFI207 constructs. NIH3T3 cells were transfected with full length or mutant IFI207 expression constructs. Cells were fixed and stained with antibodies directed against expression construct tags. Images were acquired by fluorescence microscopy. Graph below shows the quantification of cells from multiple fields.

**Fig. S8.**
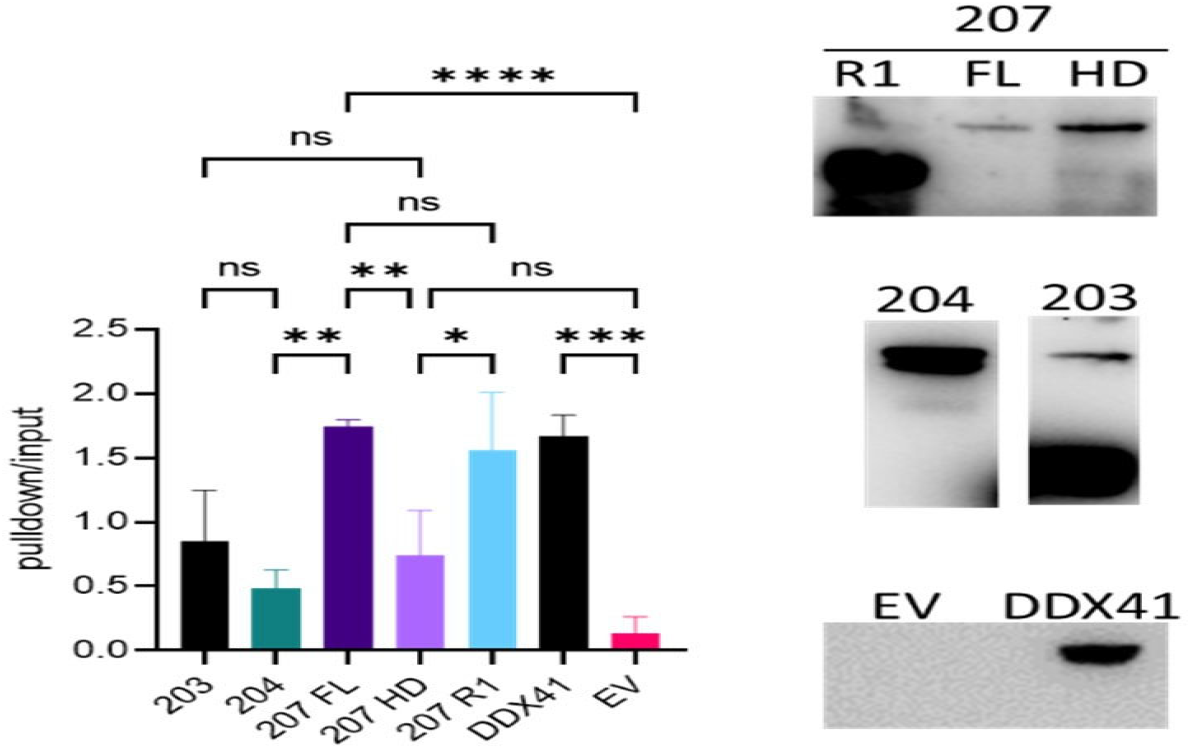
IFI207 R1 but not HD mutants bind reverse transcripts. DNA pulldown assays with extracts from cells transfected with the indicated constructs. Anti-HA was used for the IFI constructs, and anti-myc was used for DDX41. Strong stop early reverse transcript primers were used to carry out qPCR. Average of 3 independent experiments with S.D. *, *P* ≤ 0.02; **, *P* ≤ 0.008; ***, *P* ≤ 0.0002; ****, *P* ≤ 0.0001 (two-tailed T test). Right panel: Extracts from 293T-mCAT cells transiently transfected with the indicated expression vectors were immunoprecipitated with anti-HA antibodies and analyzed on western blots. Shown are the results of a single experiment (representative of 3 independent experiments).

**Fig. S9.**
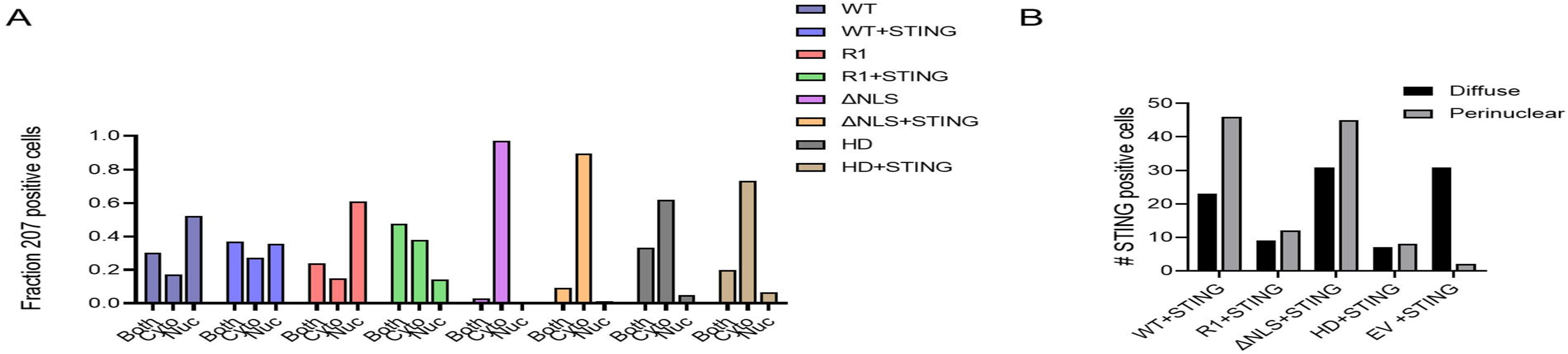
Subcellular localization of STING and IFI207. (**A**) Localization of IFI207 in the presence and absence of STING co-transfection. Single transfections are the same data presented in Fig. S7. (**B**) Localization of STING in the cytoplasm of cells co-transfected with the different IFI207 mutants.

**Table S1.**
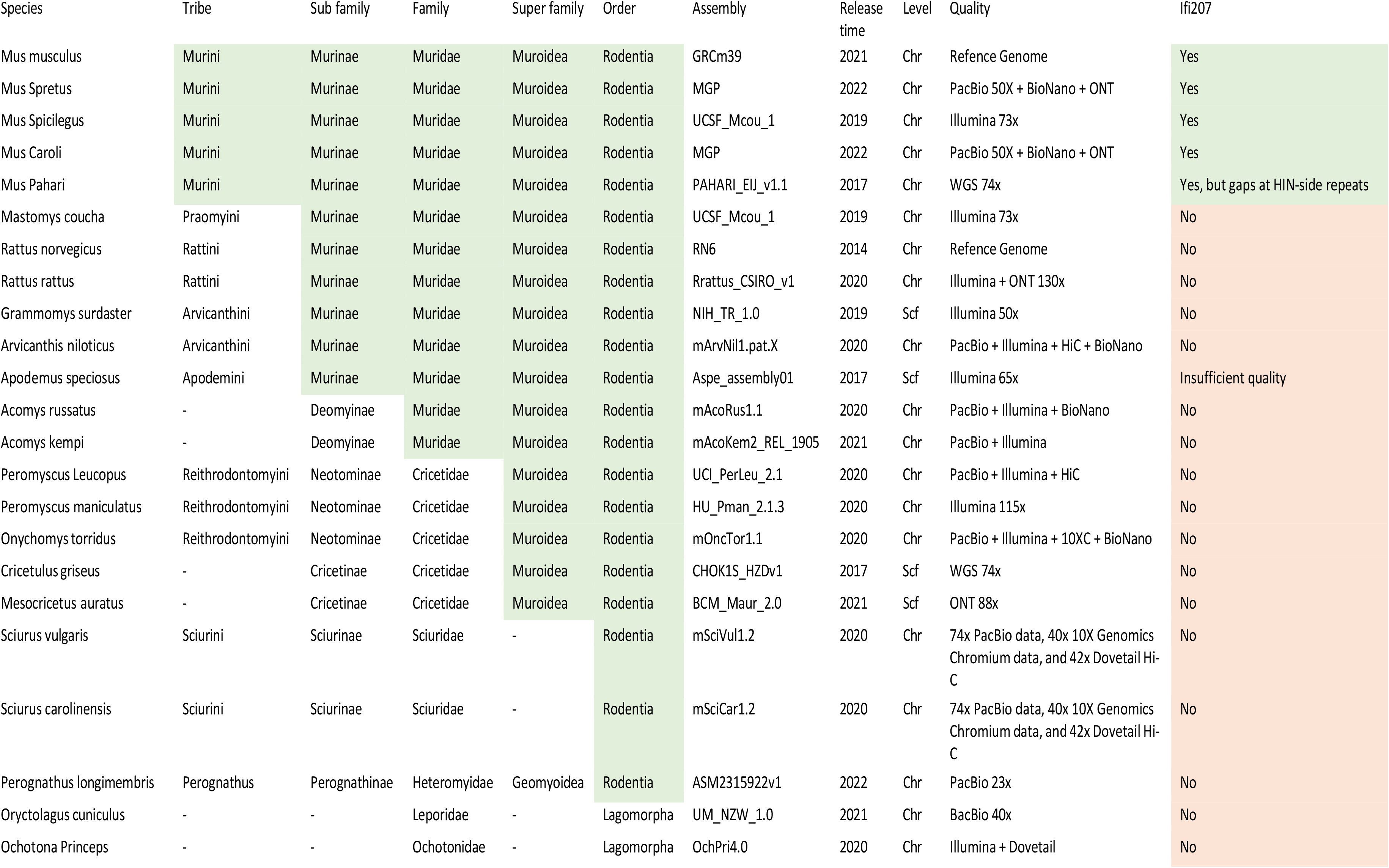
Species analyzed for the presence of Ifi207.

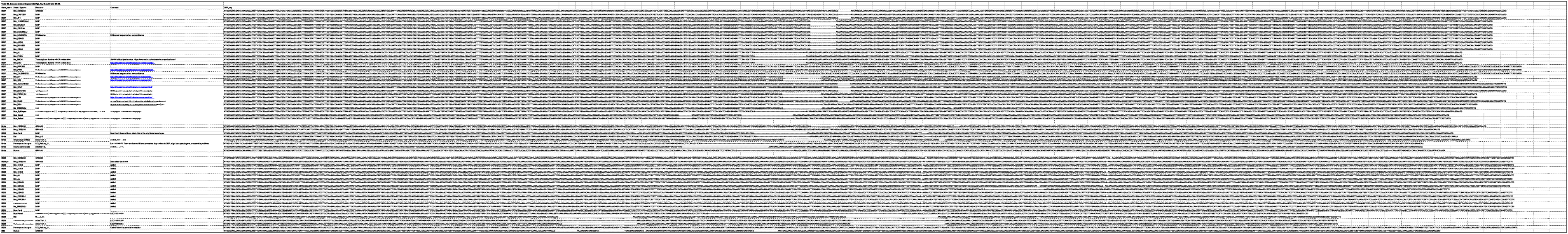

**Table S3.**
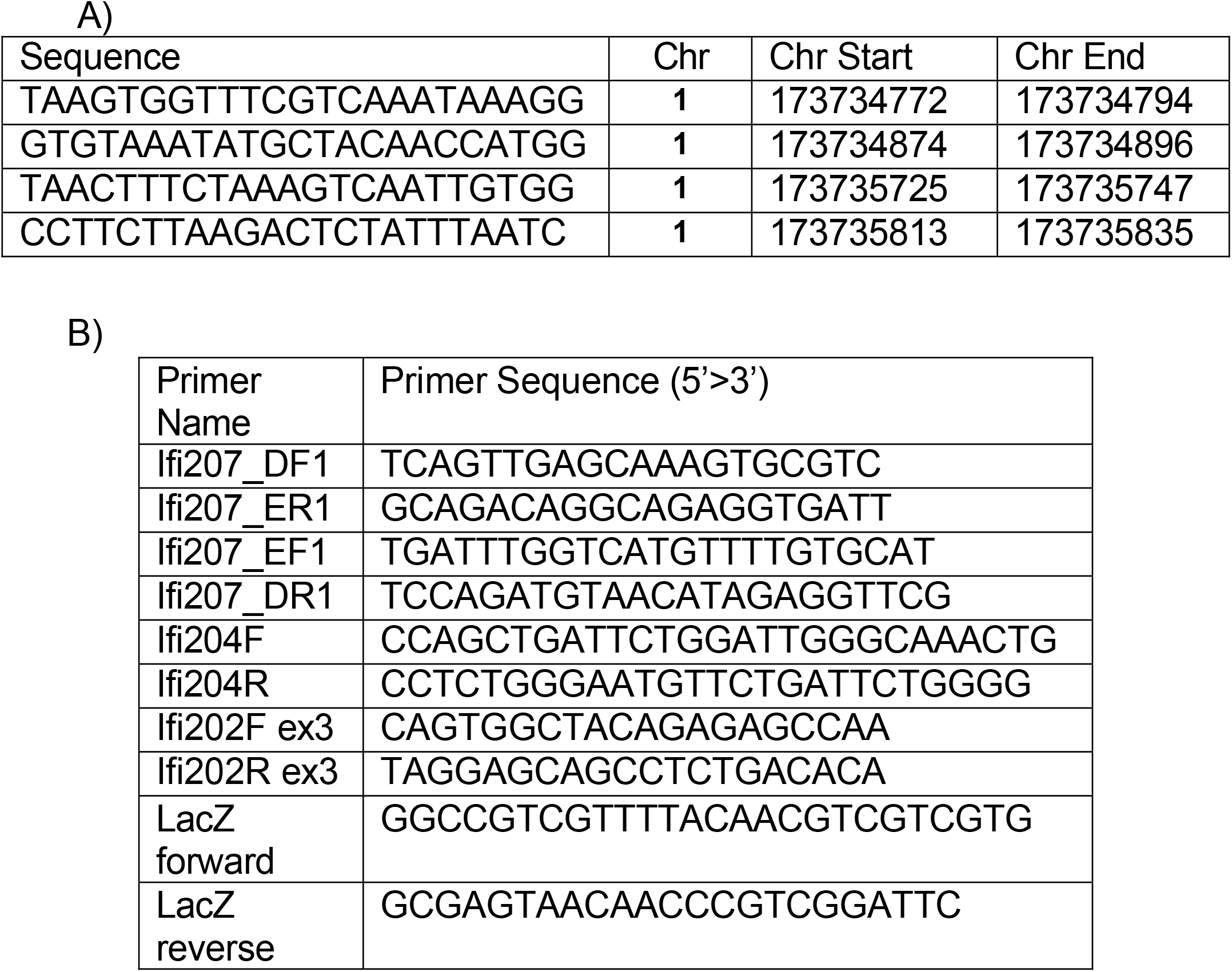
Guide RNAs used to generate IFI207 knockouts (A) and (B) PCR primers used to genotype IFI207, IFI204 and IFI202 knockout mice. IFI202 knockout mice were genotyped by the presence of LacZ sequences and the absence of exon 3.

**Table S4.** Genbank accession numbers for sequences.

| Mouse Strain | Source | Description | GenBank accession # |
| --- | --- | --- | --- |
| DBA/2J | cDNA | Full length cDNA | ON550581 |
| 129P2/OlaHsd | cDNA | Full length cDNA | ON550582 |
| C57BL/6N | cDNA | Repeat region | ON550583 |
| DBA2J | cDNA | Repeat region | ON550584 |
| Aim2<sup>-/-</sup> (B6.129P2-Aim2<sup>Gt</sup>(CSG445)Byg/J) | cDNA | Repeat region | ON550585 |
| 129P2/OlaHsd | cDNA | Repeat region | ON550586 |
| BALB/cJ | cDNA | Repeat region | ON550587 |
| M. M. castaneus | cDNA | Repeat region | ON550588 |

**Table S5.**
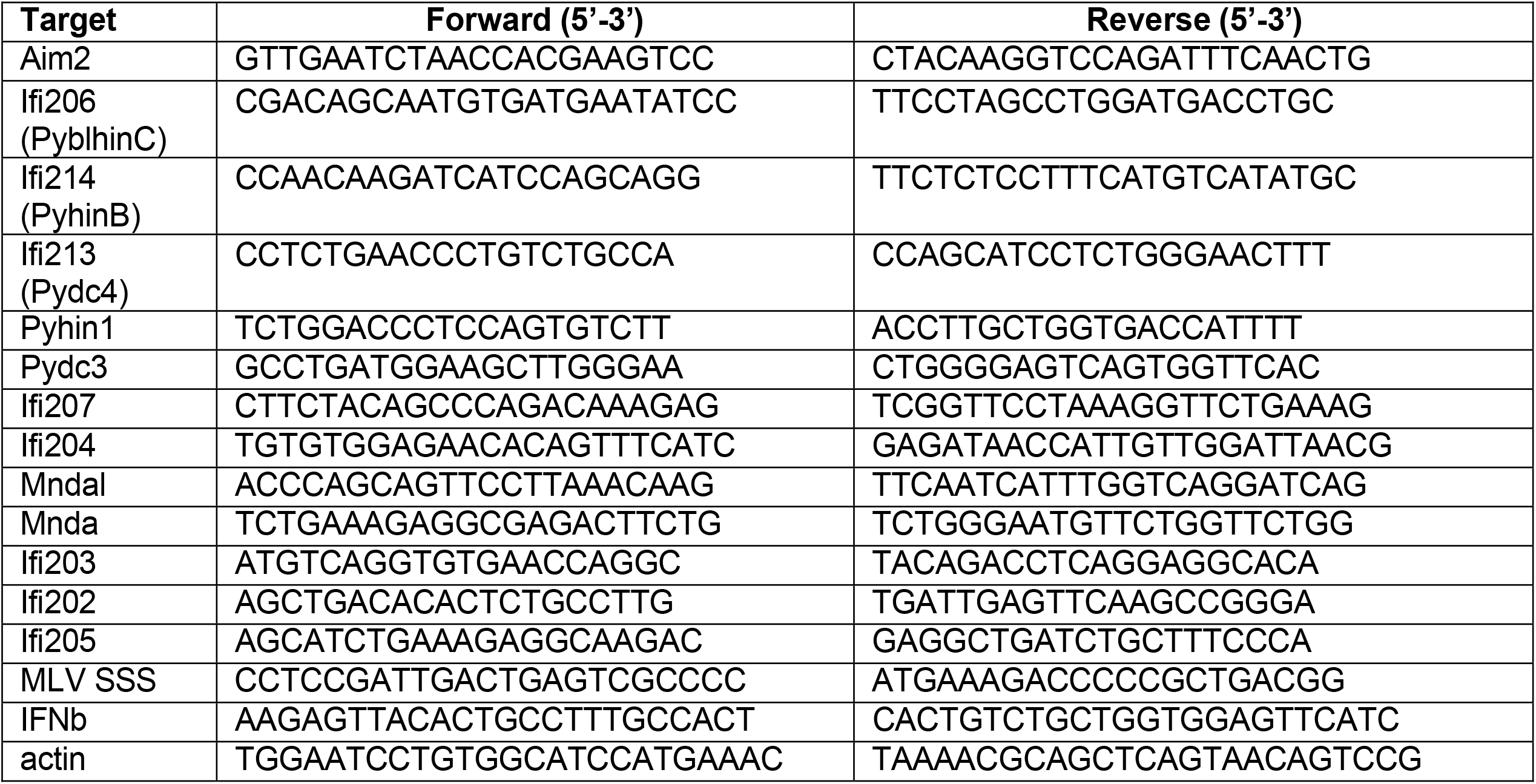
Primers used for expression analysis (Fig. 2) and MLV reverse transcript pulldowns (Fig. S8).

**Table S6.**
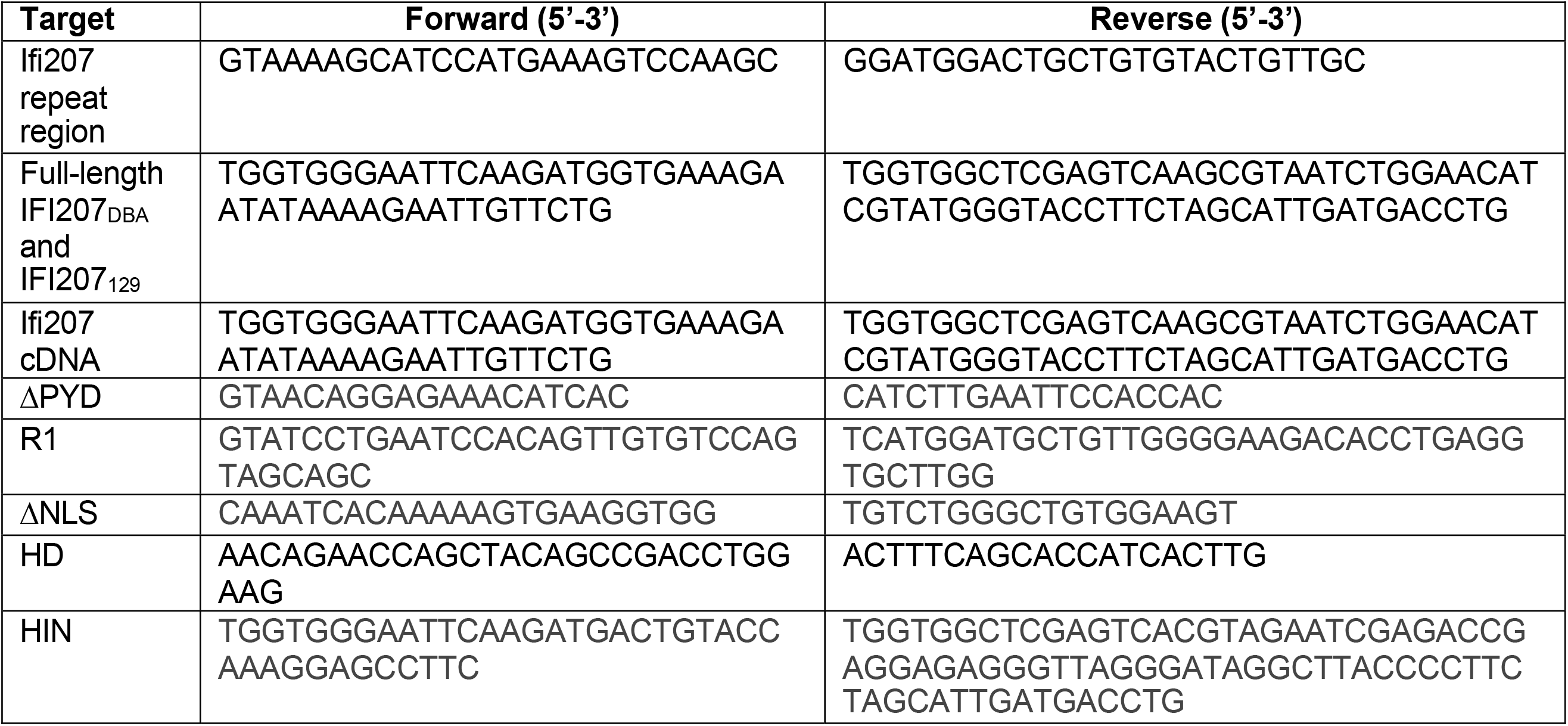
Sequencing, cloning and mutagenesis primers.

